# Reconstruct a eukaryotic chromosome arm by *de novo* design and synthesis

**DOI:** 10.1101/2022.10.04.509869

**Authors:** Shuangying Jiang, Zhouqing Luo, Kang Yu, Shijun Zhao, Zelin Cai, Wenfei Yu, Hui Wang, Li Cheng, Zhenzhen Liang, Hui Gao, Marco Monti, Daniel Schindler, Linsen Huang, Cheng Zeng, Weimin Zhang, Chun Zhou, Yuanwei Tang, Tianyi Li, Yingxin Ma, Yizhi Cai, Jef D. Boeke, Junbiao Dai

## Abstract

The genome of an organism is inherited from its ancestor and keeps evolving over time, however, how much the current version could be altered remains unknown. Here, we use the left arm of chromosome XII (*chrXIIL*) as an example to probe the genome plasticity in *Saccharomyces cerevisiae*. A neochromosome was designed to harbor originally dispersed genes. The essentiality of sequences in *chrXIIL* was dissected by targeted DNA removal, chromosome truncation and random deletion. Notably, 12 genes were sufficient for survival, while 25 genes are required to retain robust fitness. Next, we demonstrated these genes could be reconstructed using synthetic regulatory sequences and recoded open-reading frames with “one-amino-acid-one-codon” strategy. Finally, we built a neochromsome, which could substitute for *chrXIIL* for cell viability, with these reconstructed genes. Our work not only highlights the high plasticity of yeast genome, but also illustrates the possibility of making functional chromosomes with completely artificial sequences.

**HIGHLIGHTS:**

1. A neochromosome was designed to facilitate the assembly of exogenous DNA for stable expression in yeast
2. The left arm of *chrXII* could be minimized to just 12 genes to maintain viability, but additional genes were required to retain robust fitness
3. Comprehensive recoding and transcriptional refactoring using artificial regulatory sequences produced a functional chromosome arm
4. A completely reconstructed neochromosome could replace the *chrXIIL* to maintain comparable fitness

## INTRODUCTION

*Mycoplasma genitalium* was regarded as the bacteria with the smallest genome that can grow in axenic culture (Fraser et al., 1995; Glass et al., 2006). In 2016, the record was broken after the creation of JCVI-syn3.0 with 531 kbp in genome size, reduced from over 1.0 Mbp in original *Mycoplasma mycoides* (Hutchison et al., 2016). In this genome, only 473 genes, including the set of essential and quasi-essential genes, were retained, producing a near-minimal bacteria cell (Hutchison et al., 2016; Pelletier et al., 2021). On the other hand, the 4.6 Mbp *Escherichia coli* MG1655 genome encodes 4,434 genes (Blattner et al., 1997). Through rounds of deletion, the MG1655 genome was reduced by up to 15%, of which 743 genes including mobile DNAs were deleted (Posfai et al., 2006). This genome reduced strain is reported to not only preserve good growth profiles and protein production but also have beneficial properties including high electroporation efficiency and high stability of recombinant plasmids.

Compared to prokaryotes, genome reduction studies in eukaryotes are limited. As a model eukaryotic genome, the genome of *Saccharomyces cerevisiae (S. cerevisiae)* is 12 Mbp in length and encodes about 6000 genes (Goffeau et al., 1996). Based on chromosome-splitting and losing techniques, it has been reported that the 5% reduction of the *S. cerevisiae* genome could improve productivity of ethanol and glycerol (Murakami et al., 2007). Since 2006, a group of researchers have devoted to construct a designer version of the yeast genome (known as Sc2.0), in which the yeast genome size will be reduced up about 8% through deletion of the retrotransposon related sequences, introns and subtelomeric repeat sequences (Richardson et al., 2017). In the Sc2.0 genome, loxPsym sites were inserted 3 bp downstream of stop codon of every nonessential gene to build a system called SCRaMbLE (Synthetic Chromosome Rearrangement and Modification by LoxPsym-mediated Evolution)(Dymond et al., 2011). Upon activation of Cre recombinase, these loxPsym sites will mediate random inversion, deletion and duplication of genomic sequences (Jiang et al., 2020b). Recently, we developed an iterative SCRaMbLE-based genome compaction (SGC) strategy to compact the left arm of synthetic chromosome XII (*synXIIL*), revealing that at least 39 of the 65 nonessential genes in *synXIIL* are not required for cell viability at 30°C in rich medium (Luo et al., 2021).

Both coding and regulatory gene sequences are essential, and define the basic functional unit of a genome. Due to codon degeneracy and usage bias, different nucleotide sequences encoding the same amino acid sequence might have different functional impacts (Komar, 2016). The effects of synonymous recoding are difficult to predict. Only in *E. coli* has genome-wide synonymous codon compression been successfully applied through genome editing (Isaacs et al., 2011; Lajoie et al., 2013) or whole genome synthesis (Fredens et al., 2019). Permissive and restrictive synonymous recoding schemes are largely unexplored in eukaryotes. While the “genetic code” determines a protein’s amino acid sequence, these noncoding regions determine when and where these proteins are produced according to various “gene regulatory codes”. Many functional elements such as enhancers, transcription factor binding sites, the TATA box, the transcription start site and poly(A) site belong to these regulatory codes. Meanwhile, synthetic promoters and terminators have been designed based on these codes and have shown varied activities (Curran et al., 2015; de Boer et al., 2020; Redden and Alper, 2015). How to match the regulatory code to gene function genome-wide is still an un-explored field in synthetic genomics.

The left arm of *chrXII* in *S. cerevisiae* (*chrXIIL*) is 150,827 bp long, containing 73 protein coding genes, 1 tRNA coding genes, 3 autonomously replicating sequences and 1 Ty1 LTR (SGD, https://www.yeastgenome.org/). Among these genes, 10 are defined as essential based on the phenotype of individual knockout mutations and their ability to support spore viability in a heterozygous diploid. Our recent work using SCRaMbLE indicated that at least half of the genes in *chrXIIL* were not necessary for cell viability (Luo et al., 2021). However, how much of the arm can be further reduced in size remains unknown. Furthermore, whether we can use a limited number of artificially “reconstructed” genes (with heterologous cis-regulatory sequences) to replace the *chrXIIL* for survival is unknown. In this study, a neochromosome was initially designed to facilitate the relocation of essential genes which are dispersed throughout *chrXIIL*. Using a partially synthetic chromosome XII, we systematically probed sequence essentiality in *chrXIIL* by targeted DNA deletion, telomere capping and random deletion using SCRaMbLE. Eventually, a series of designed neochromsomes, capable of substituting for *chrXIIL* for cell viability were constructed.

## RESULTS

### Design of a neochromosome as a flexible carrier of exogenous DNA

Linear artificial chromosomes containing structural elements for segregation, replication and stability (*i*.*e*., centromere, autonomous replication sequences and telomeres) have been reported to behave like the native ones (Dani and Zakian, 1983; Murray and Szostak, 1983). However, *de novo* construction of a linear chromosome with capability for future applications remains limited. To produce a maximally useful resource, we designed a neochromosome with unique features allowing it to 1) be stably propagated and inherited without any need for selection, 2) accept exogenous DNA fragments with high efficiency, and 3) be assembled easily *in vivo*. The overall construct design is shown in Figure 1A.

**Figure 1.**
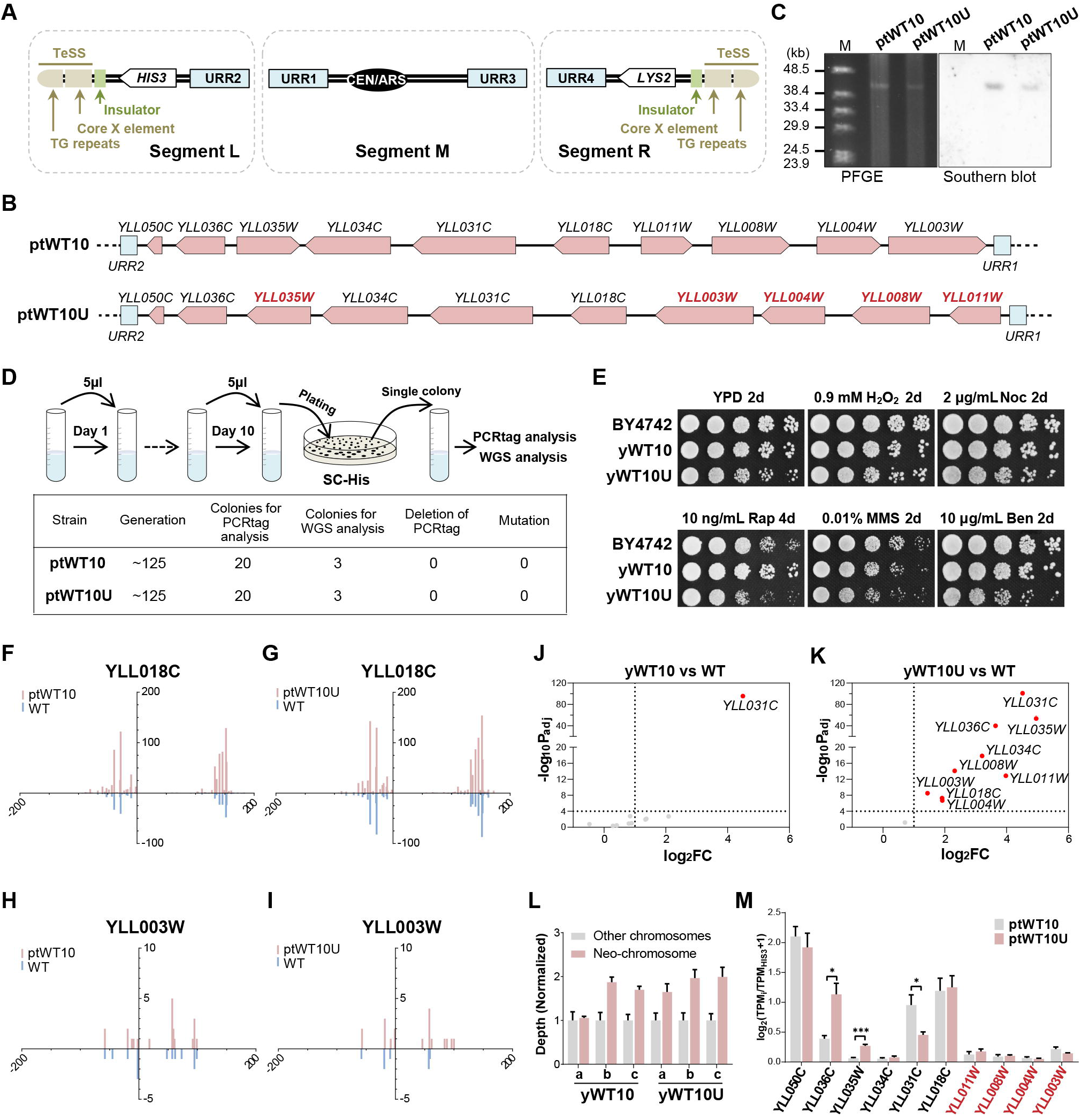
A designer neochromosome to harbor essentials genes from the left arm of *chrXII*. (A) The design of neochromosome. Marker genes, white; TeSS, brown; Insulators, green; URRs, blue. (B) Arrangements of the ten essential genes (pink) in ptWT10 and ptWT10U. Arrows point to the direction of transcription. The genes with changed transcription direction on ptWT10U were labeled by red. (C) PFGE and southern blot analysis of the assembled neochromosomes. M: λ DNA-Mono Cut Mix. (D) Stability test of neochromosomes. BY4742 cells containing ptWT10 or ptWT10U were cultured in SC-His medium for about 125 generations before PCR analysis and whole genome sequencing. (E) Phenotypic analysis of strains on different media. H_2_O_2_: hydrogen peroxide; Noc: nocodazole; MMS: methyl methanesulfonate; Rap: rapamycin; Ben: Benomyl. (F-I) Full-length transcriptome analysis of BY4742 with ptWT10 or ptWT10U neochromosome. The transcripts on neochromosome (pink) and native transcripts (blue) were mirrored. In x-axis, negative number represents distance upstream of ATG and positive value indicates distance downstream of the stop codon. The Y-axis represents the counts of detected transcripts. (J-K) Transcription analysis of the ten essential genes in yWT10 or yWT10U. Differential expression in this study was defined as |Log_2_ (Fold change)| > 1 and -log_10_ (Adjusted p-value) >4. Dashed lines, the threshold defined above. Red, differentially expressed essential genes on neochromosomes. (L) The copy number of neochromosomes in yWT10 or yWT10U. a,b,c are three independent clones. In this study, the error bar is the SD. (M) Normalized expression of genes on ptWT10 and ptWT10U. Expression of each gene was normalized to *HIS3* on neochromosomes. *P<0.05, **P<0.01, ***P<0.001, ****P<0.0001.

The telomere seed sequences (TeSS), that is, the telomeric TG repeat plus the Core X element, which have been constructed and functional tested previously, served as the telomeres (Richardson et al., 2017). To ensure replication and segregation of the neochromosome, the centromere sequence from Chromosome II and an origin of replication (ARS208) were placed in the middle. It is well known that genes which are placed near telomere, will be silenced - a phenotype called telomere position effect (TPE). Therefore, to prevent TPE, a short insulator sequence (10 tandem repeats of TTAGGG) was put adjacent to TeSS to block the spreading of heterochromatin (Fourel et al., 1999). In addition, to allow robust integration of exogenous DNA with high efficiency and specificity, two 500bp random sequences, designated as the universal recombination regions (URRs) were also included in each chromosomal arm (Guo et al., 2015). Finally, two auxotrophic markers, *HIS3* and *LYS2* were inserted into the left and right arm respectively, adjacent to the URRs to facilitate the selection of the assembled neochromosome. The entire structural elements of the neochromosome were divided into three parts, fragments L, M and R, which could be assembled together in yeast by homologous recombination.

### Construction of neochromosomes carrying essential genes from *chrXIIL*

We used the designed neochromosome to carry the 10 known essential genes from *chrXIIL*. We positioned these genes in two different formats (Figure 1B): 1) in the same relative chromosomal position and transcriptional orientation as on the native chromosome (designated as ptWT10), and 2) unifying the transcriptional direction to point towards the telomere (ptWT10U). To facilitate assembly, four genes *YLL003W, YLL004W, YLL008W, YLL011W* were flipped together, which lead to the change of their relative position (Figure 1B). In addition, since *YLL035W* and *YLL036C* share a bi-directional promoter, we arbitrarily chose p*CYC1* to drive the expression of *YLL035W* and *YLL036C* in ptWT10U (Liang et al., 2022). The sequences of ptWT10 and ptWT10U are listed in Table S1.

To construct the neochromosomes, we developed a one-step assembly method named Transformation-assisted Linear Chromosome Construction (TLCC, Fig S1A). In TLCC, the three structural fragments, i.e., fragment L, M and R, together with PCR-amplified essential-gene fragments are co-transformed into a yeast strain. Subsequently, cells containing assembled chromosomes were selected in a medium lacking histidine and lysine, randomly isolated and confirmed by PCR (Figure S1B-S1E). Next, the strains were analyzed by PFGE followed by Southern Blot. As shown in Figure 1C, a band at the size of around 40 kb was detected in both PFGE and southern Blot, indicating the successful construction of the neochromosome. Finally, whole genome sequencing (WGS) was performed which revealed that both ptWT10 and ptWT10U are present in the cells with sequences as designed (Figure S1F-1G). However, we observed several single nucleotide substitutions and insertions in the neochromosome (Figure S1H-1I) which do not affect the function of each essential gene (see below), and therefore, no further correction was performed. Interestingly, we found that the telomeric TG repeats, in both neochromosomes, significantly expanded (Figure S1J-1K), indicating the TeSS end grew a new telomere successfully, as designed (Jiang et al., 2020a; Richardson et al., 2017).

Length has been reported to be a critical factor affecting the stability of the linear mini-chromosomes during mitosis (Murray and Szostak, 1983). Since the neochromosomes are relatively short (less than 50 kb) and were built from synthesized DNA fragments, we first tested their stability in cells (Annaluru et al., 2014). As shown in Figure 1D, we inoculated the strains in liquid medium and subcultured the cells by 1000-fold dilution every day for ten consecutive days, accounting for about 125 mitotic generations. The cells were analyzed by both PCR analysis and whole genome sequencing. For three independent clones tested, we found that none of them had changes in PCRtags or nucleotide sequences. These results demonstrate that the linear neochromosomes are stable in the cells.

### Relocation of genes to the neochromosome has certain effects on their functions

To test whether genes relocated to neochromosomes function equivalently to those in native loci, we constructed strains with ptWT10 or ptWT10U in which the original genomic copy of these essential genes were all deleted (designated as yWT10 and yWT10U). We found that both strains grew normally (Figure 1E). Interestingly, although yWT10 showed indistinguishable growth from that of wild type under various stress conditions, yWT10U exhibited certain growth defects (Figure 1E). These results suggest that the general function of these essential genes is largely unaffected after relocation and the orientation of transcription seems not to be crucial.

Since we restructured the position of these essential genes by placing them side by side, especially in ptWT10U, it might lead to interference between adjacent transcription units (Brooks et al., 2022). Therefore, we examined whether there were any transcriptional abnormalities in transcription initiation and termination sites in each gene in the two neochromosomes. From the isoform-sequencing (Iso-seq) results of BY4742 with either the ptWT10 or ptWT10U neochromosome, the full-length transcripts of the 10 genes on the neochromosomes, were identified using specific PCRtags (except for *YLL050C*, which is too short to contain PCRTags) (Zhang et al., 2017). For genes with the same transcription direction, such as *YLL018C*, the transcription start sites (TSS) and termination sites (TES) on the neochromosomes are similar to those in the native transcriptome (Figure 1F-1G, S2A-B). For genes with altered directions on the neochromosomes, such as *YLL003W*, they were also transcribed similarly to the start and termination sites (Figure 1H-I, S2C-E). For *YLL035W* and *YLL036C*, the TES and TSS showed a similar pattern between ptWT10 and the native genome locus, while the promoter and terminator of *CYC1* used in ptWT10U changed the transcription pattern as expected (Figure S2F-H). In addition, we failed to observe abnormal readthrough between two adjacent genes. These results indicated that the sequences we used for each gene were sufficient to regulate the transcription initiation and termination processes.

To examine whether the transcription level of genes was altered when relocated to the neochromosome, the expression of the ten genes in yWT10 and yWT10U was analyzed by RNA-seq (Figure 1J-K). We found that in yWT10, all 10 essential genes except for *YLL031C* were transcribed at similar levels as their genomic counterparts in BY4742. Unexpected, 9 out of the 10 essential genes, including the two with altered transcription regulatory sequences, were overexpressed in yWT10U (Figure 1K and see below). Finally, consistent with the above Iso-seq results, the transcripts of most genes identified on the neochromosomes showed clear boundaries between different transcription units (Figure S2I).

### The neochromosomes exist in cells with variable copy number

The elevated expression of genes in yWT10U lead us to ask whether the copy number of neochromosome had changed. We looked into the sequence data and divided the reads into two parts: the linear neochromosome and other remaining chromosomes. The mean of the sequencing depth of a 1 kb-sized bin for all the other chromosomes was normalized to 1 and the ratio of the mean depth of the neochromosome to all other native chromosomes was calculated as the mean copy number of the neochromosome per haploid genome. As shown in Figure 1L, we found the neochromosomes exist in cells with 1 to 2 copies.

To eliminate the effects of DNA copy number, we normalized the expression level of each gene on the neochromosome to that of *HIS3*, which is located on the left arms of the neochromosomes. As shown in Figure 1M, three genes, including the two with changed promoters and *YLL031C*, showed obvious different transcription levels between ptWT10 and ptWT10U, while none of the four genes with altered directions, namely *YLL011W, YLL008W, YLL004W* and *YLL003W*, showed different transcription levels. These results suggest that the sequence, rather than the orientation of transcription, plays important roles in gene regulation.

Together, these results suggest that neochromosomes constructed using artificially designed structural elements can be carriers for essential genes, even if they are refactored. Similar to the discovery that genetic organization is important but not critical for prokaryotes (Hutchison et al., 2016), our data also supports the conclusion that although the eukaryotic gene arrangement impinges upon expression and strain fitness, it is not a major determinant for survival, highlighting the plasticity of gene arrangement in eukaryotic genome.

### Only 12 genes are required for viability within the left arm of *chrXII*

Previously, we found over half of the nonessential genes in *synXIIL* could be deleted by SCRaMbLE (Luo et al., 2021). However, further compaction attempts using the same method failed, which may be partially due to the extremely slow growth of the final strain. To probe the minimal gene set to support cell viability, two additional strategies were carried out here.

At first, we systematically examined the essentiality of genome blocks in the left arm of *chrXII*. Six genome blocks flanking essential genes that have not been fully deleted by SCRaMbLE in our previous study (Luo et al., 2021) were knocked out individually by CRISPR/Cas9 technology (Figure S3A). Notably, all deletions generated viable strains, despite that two of them exhibited growth defects (Figure S3B). These results suggested that all these non-essential regions could be removed, respectively and there is no essential “dark matter” in these regions.

Next, a method called Tel-cap, which is similar to previous technique for the replacement of telomere (Ray and Runge, 1999), was adopted to truncate the chromosome arm piece by piece (Figure 2A). In this method, a fragment containing a universal telomere, a marker gene and a homologous region is transformed into yeast to create a new telomere at the left end of *chrXII* by homologous recombination. Depending on the location of the homologous region (HR I-IV, Figure S3A), the left-most arm up to this region will be deleted. Using Tel-cap, we systematically remove the left arm of *chrXII* with quarterly increments to a maximum of the entire arm in a heterozygous diploid strain (Region I-IV, Figure 2B). Due to deletion of essential genes, the diploid strains with an empty neochromosome backbone (Neo0) produced only two viable spores upon sporulation (Figure 2B). In contrast, the presence of ptWT10 rescued the lethality of the two spores in three strains, except for the one in which the entire *chrXIIL* was deleted (*chrXIIL*Δ, Figure 2B). These results indicate that only 10 essential genes are insufficient to substitute for *chrXIIL* and that additional gene(s) in region IV are required.

**Figure 2.**
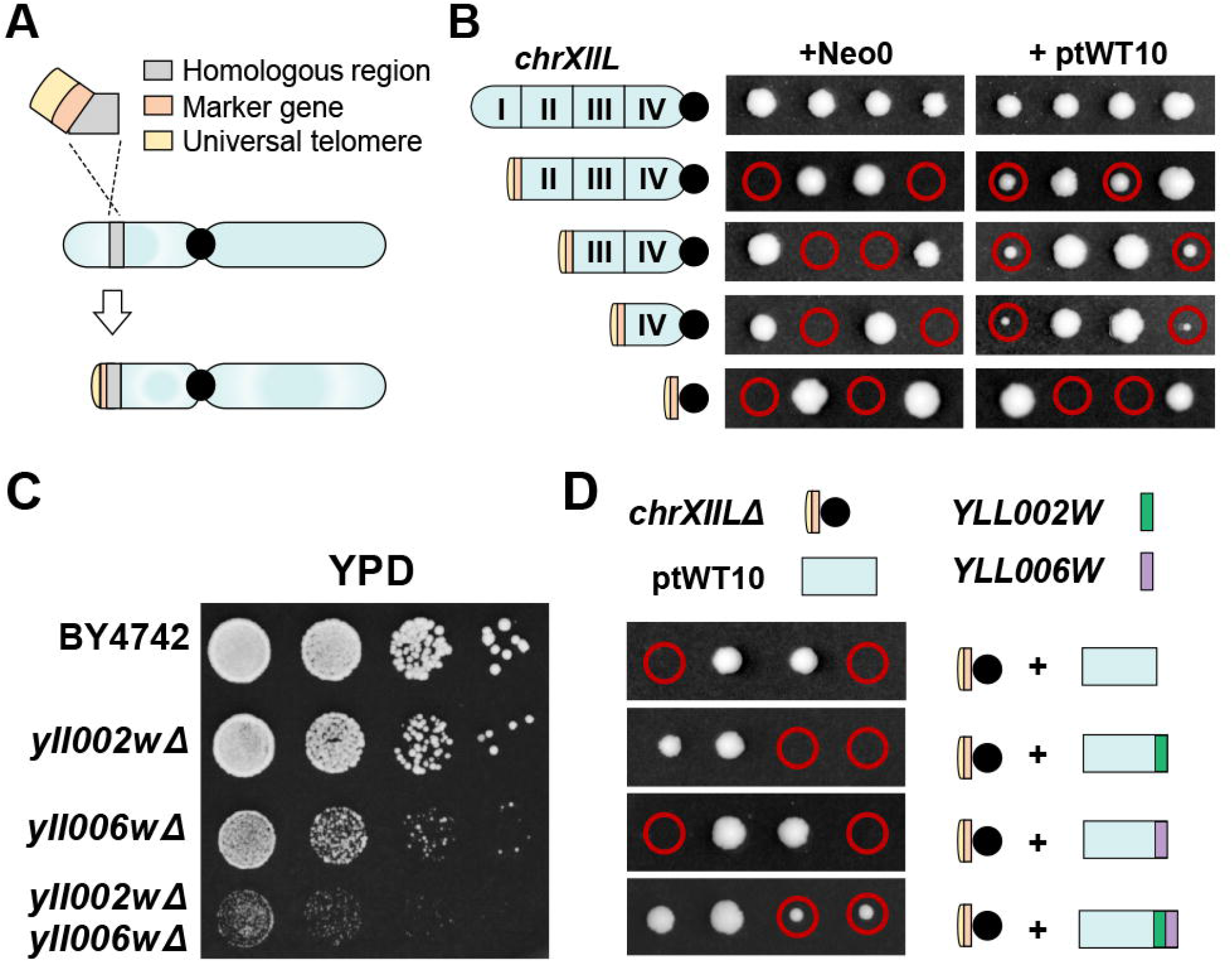
Only 12 genes are sufficient for viability within *chrXIIL*. (A) The schematic diagram of Tel-cap method for chromosomal truncation. (B) Tetrad analysis of heterozygous diploid strains with different *chrXIIL* truncations. (C) Fitness tests of strains carrying *yll002w* and *yll006w* deletion. (D) Tetrad analysis of heterozygous diploid strains with one of its *chrXIIL* removed. *YLL002W* (green) and *YLL006W* (purple) were integrated into the neochromosomes respectively or together. Red circle, the spores containing *chrXIIL*Δ and the neochromosome.

After careful examination of genes in region IV (region between HR III and HR IV, Figure S3A), we focused our attention on two genes, *YLL002W* and *YLL006W*, which are known to be required for cell growth (Driscoll et al., 2007; Han et al., 2007; Kornmann et al., 2009). As shown in Figure 2C, deletion of each gene respectively impairs cell fitness and the double mutant leads to a severe growth defect. Therefore, we constructed three versions of neochromosomes containing the 10 essential genes plus either *YLL002W, YLL006W*, or both. Excitingly, we found only the strain containing ptWT10 plus both *YLL002W* and *YLL006W* generated four viable spores, suggesting that a neochromosome containing just the 12 genes is sufficient to substitute *chrXIIL* for yeast viability. We named the neochromosome with 12 genes as ptWT12 and subsequently confirmed its identity by sequencing (Figure S3C-3D, Table S1). The yeast strain carrying ptWT12 and *chrXIIL*Δ was named as yWT12.

### Additional genes are needed to restore normal cell growth

Although yWT12 is viable, it grew poorly even in rich medium (Figure 2D). We hypothesized that adding back additional non-essential genes from *chrXIIL* might be able to recover the cell fitness. Therefore, we analyzed all genes in *chrXIIL* to identify potential candidates using the following criteria. Firstly, the seven genes with the most reported genetic interactions (>250) were selected (Table S2). Secondly, other genes which were both reported to cause decreased vegetative growth rates or respiratory growth defects when deleted (marked with red lines) and retained in ZLY348 (marked with green dots), a strain with compacted *chrXIIL* but retaining wild type-like growth (Luo et al., 2021). By this means, we identified a set of 13 genes that improved fitness (Figure 3A and Table S2). These genes, including their regulatory sequences, were amplified from wild type genome, added into the neochromosome and generated ptWT25 and the strain carrying ptWT25 and *chrXIIL*Δ (yWT25, Figure 3A-B, Table S1). In addition to ptWT25, we also constructed an intermediate neochromosome containing 18 genes, called ptWT18 (Figure 3A-B and Table S1). The corresponding strains were designated as yWT18. The successful construction of these neochromosomes was verified by sequencing (Figure S3 E-I). The fitness of these strains was evaluated under various stress conditions. As shown in Fig 3B, the introduce of additional genes obviously improved the growth of cells on rich medium and under the chemical stresses of MMS and rapamycin. All of the strains showed no obvious sensitivity to the oxidative stress (H_2_O_2_) or the anti-microtubule drugs (benomyl and nocodazole) used.

**Figure 3.**
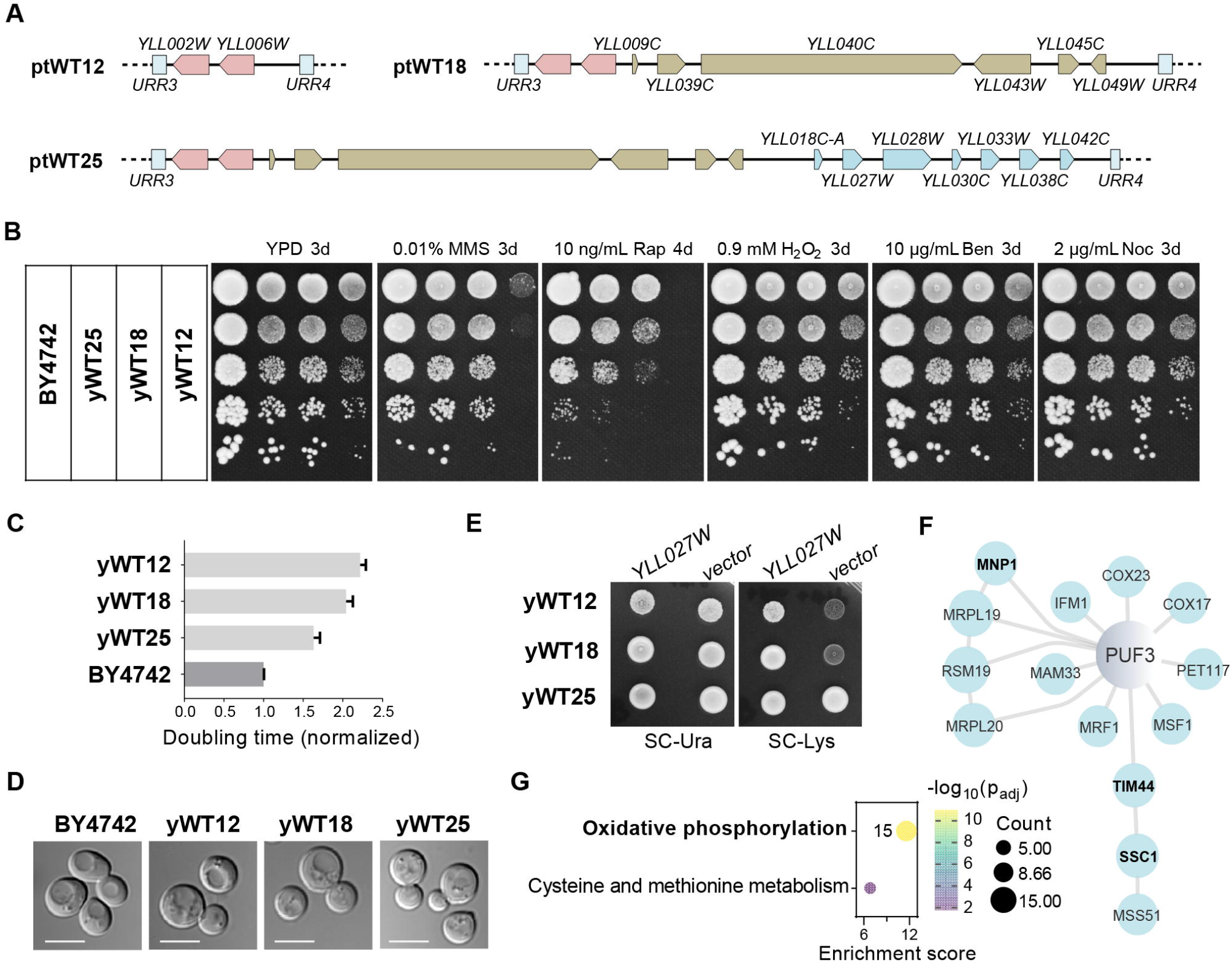
Restoration of cell fitness using a simplified gene set. (A) Schematic diagrams of additional genes in the three neochromosomes besides the essential genes. (B) Fitness analysis of strains carrying assembled neochromosomes under various conditions. (C) Doubling times of strains carrying corresponding neochromosome in YPD. The mean doubling time of BY4742 was set to 1.0. (D) Cell morphology of indicated strains. Scale bars, 5 μm. (E) Lysine auxotrophy due to *yll027w deletion. YLL027W* is expressed in a centromeric plasmid under its native promoter and terminator. (F) The interaction map between *PUF3* and genes up regulated in yWT25. The network was generated with the string database (Szklarczyk et al., 2021) (experiment and database, cutoff score is 0.7). Node size, connection number. (G) KEGG pathway enrichment analysis of down-regulated genes in yWT25.Threshold, Adjusted p-value<0.05. Bubble size, the count of genes; the bubble color, -log_10_ (Adjusted p-value).

Next, we measured the growth rates of these strains. As expected, the doubling time of yWT12 was lengthened to over twice that of wild type. The doubling time was gradually reduced for yWT18 and yWT25. However, even yWT25 grew much slower than BY4742 (Figure 3C). In addition, we analyzed whether there were morphological changes among these strains. As shown in Figure 3D, the cells all looked similar. Interestingly, neither yWT12 nor yWT18 could grow on the synthetic medium without lysine while yWT25 do (Figure 3E). This phenotype may result from the removal and subsequent re-incorporation of *YLL027W* (Hanway et al., 2002; Jensen and Culotta, 2000). Consistent with this hypothesis, expression of *YLL027W* in yWT12 or yWT18 did recover growth on the selective medium (Figure 3E).

To reveal possible reasons for the slow growth of yWT25, we compared the transcriptional profile of yWT25 to wild type. After examination of the 140 differentially expressed genes (Table S3), we found 25% (14/52) genes with increased expression are involved in the interaction network with *PUF3* (*YLL013C*, Figure 3F), which regulates gene expression through its roles in the decay of mRNA targets (Miller et al., 2014; Olivas and Parker, 2000), and three genes, namely *TIM44, SSC1* and *MNP1*, have been reported to showed decreased rate of vegetative growth when over-expressed (Koike et al., 2018; Sopko et al., 2006; Yoshikawa et al., 2011). As for the 88 down-regulated genes, many are enriched in the oxidative phosphorylation pathway which was related to *YLL041C* (*SDH2*) (Fig 3G and S3J-K). Together, these results suggest that although only 12 genes are required to *chrXIIL* for cell survival, additional genes are needed to maintain a relative robust growth. And the principles used in this study are useful to identify critical genes for strain fitness.

### Reconstruct transcription units using exogenous artificial sequences

Given that the native genes could be relocated to neochromosome with little effects on function, we next asked if both the coding and regulatory sequences are able to be reconstructed with completely synthetic ones. Previously, it has been shown that *E. coli* with only 61 codons survives (Fredens et al., 2019) and yeast strains lacking TAG codon in one chromosome grows indistinguishable from that of wild type (Annaluru et al., 2014; Jiang et al., 2020a). In addition, synthetic promoters and terminators have been designed and showed varied activities (Curran et al., 2015; de Boer et al., 2020; Redden and Alper, 2015). However, to what extent codons can be simplified and moreover whether an entire chromosomal arm could be functionally reconstructed remains to be tested.

In eukaryotes, a gene typically consists of three parts, *i*.*e*., coding sequences (CDS), promoter elements (PRO) and terminator/polyadenylation sequences (TER). For the 25 genes in ptWT25, we systematically reconstructed them using the following principles: For CDS, the optimized DNA sequences were generated using GeneDesign software (Richardson et al., 2010), which leading a radical codon compression scheme, in which only one codon is used for each amino acid (Liang et al., 2022). For PRO and TER, 44 PRO and 28 TER were chosen from previous reports (Curran et al., 2015; de Boer et al., 2020; Redden and Alper, 2015). Each part was synthesized and cloned into the YeastFab vectors we developed previously (Guo et al., 2015). To obtain functional combinations of PRO, CDS and TER for each gene, we mixed the PRO-, TER-containing plasmids together with a particular CDS-containing plasmid to assemble a pool of TUs, which were subsequently transformed into the corresponding haploid deletion mutant (Figure 4A). For essential genes, plasmid shuffling was performed to identify the viable clones. For non-essential genes, fitness change under a particular growth condition was identified for each knockout strain and clones which are able to restore growth to that of wild type were collected. The plasmids were extracted, transformed into *E. coli* and subsequently sequenced to obtain the identity of PRO and TER. Different combinations of PRO and TER for each gene were re-tested to confirm their functionality, either by tetrad-based analysis of reconstructed essential genes for the ability to support cell survival (Figure 4B) or by serial-dilution analysis (Figure 4C, Figure S4 and Table S4). By these means, we were able to reconstitute 21 TUs using completely synthetic parts except *YLL031C, YLL028W, YLL003W* and *YLL039C*. For both *YLL031C and YLL028W*, we found the recoded CDS failed to complement the function of native CDS, either lost cell viability or failed to grow under stress conditions. In addition, we discovered that *YLL003W*, a gene required for G2/M transition, lost function when constitutively expressed. To obtain a functional TU, the PRO of *SWI6*, a transcription factor activating transcription during G1/S transition, was employed which, luckily, could restore the function of *YLL003W* (Liang et al., 2022). As for *YLL039C (UBI4)*, we failed to assemble the recoded gene, potentially due to its repetitive nature since it encodes 5 head-to-tail ubiquitin repeats within CDS (Finley et al., 1987).

**Figure 4.**
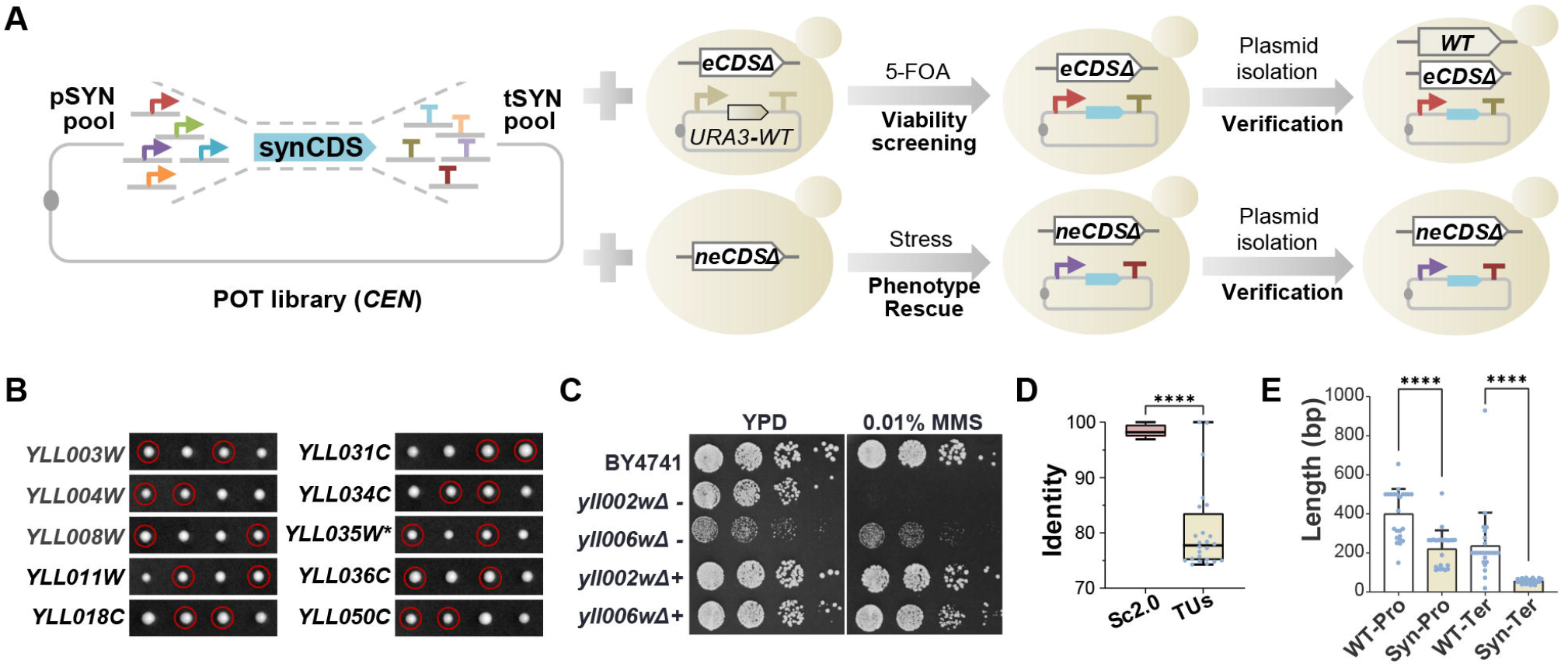
Reconstruction of transcription units. (A) Strategy to identify functional synthetic promoters and terminators to support both essential and nonessential genes. eCDS, CDS of essential gene. neCDS, CDS of nonessential gene. (B) Functional analysis of reconstructed genes by tetrad analysis. The corresponding heterozygous diploid containing a reconstructed TU (rTU) was sporulated and dissected. One representative tetrad was shown for each strain. For *YLL035W*, the ATG codon was mutated (marked with *) to avoid impact on the expression of *YLL036W*. Red circle, the spores containing disrupted native gene and reconstructed TU. (C) Functional complementation test of reconstructed *YLL002W* and *YLL006W*. + and - indicate strains with or without the reconstructed TU. (D) Comparison of sequence identity in CDS between rTUs and Sc2.0 project. (E) Length comparison of Pro and Ter between rTUs and the native ones.

Next, we compared sequence changes in the 24 TUs after reconstruction to those in wild type or Sc2.0 strain. At first, significant reduction of sequence identities in CDS was present. The Sc2.0 sequences share over 95% identity to wild type, whereas the mean identity of the reconstructed genes dropped below 80% (Figure 4D). As for the regulatory sequences, not only totally different DNA sequences were used (Table S4), the lengths of the synthetic promoters and terminators were also much shorter than the counterparts in ptWT25 (Figure 4E).

### Replacement of *chrXIIL* by a completely revamped neochromsome

Given each refactored gene is able to function similarly to the wild-type, we tested whether the combination of these genes could entirely replace *chrXIIL*. The ptSYN10 neochromosome with the 10 refactored essential genes was constructed (Figure S5A) and those essential genes were deleted from the native chromosomal locations to get the strain ySYN10 (Figure S5B). As shown in Figure S5B, ptSYN10 was able to support the viability of the ySYN10 strain, but the cells showed growth defects on either YPD or upon treatment of different drugs. RNA-seq analysis indicated that, in contrast to the neochromosomes made up of essential genes with native regulatory sequences (Figure S2I), nearly all the intergenic sequences of the synthetic genes on ptSYN10 were highly transcribed (Figure S5C). To further look into the fidelity of transcription of each gene, ptSYN10 in BY4742 was analyzed by Iso-seq (Figure S5D). Intrusive readthrough was identified into adjacent transcription units (Figure S5D). Moreover, many anti-sense transcripts appeared, especially for *syn035w* and *syn034c* (Figure S5D). These results suggested that the short synthetic terminators might not be effective at terminating transcription, and insertion of artificial DNA might bring in sequences with unexpected promoter activities.

We next built a neochromosome (ptSYN12) carrying the same 12 genes as ptWT12 but using recoded sequences (Figure S5E). Unfortunately, unlike its wild type counterpart, the ptSYN12 neochromosome was not able to substitute *chrXIIL* for survival, although it does support several cell divisions after spore germination (Figure S5F). Next, according to what we learn above using native genes, we constructed the ptSYN24 neochromosome (Figure S5G-J and Table S1), which contains 24 recoded TUs from ptWT25 except for *UBI4*. Excitingly, as shown in Figure 5B and S5K, we found ptSYN24 could support viability in spores with *chrXIIL*Δ. These strains were designated as ySYN24.

**Figure 5.**
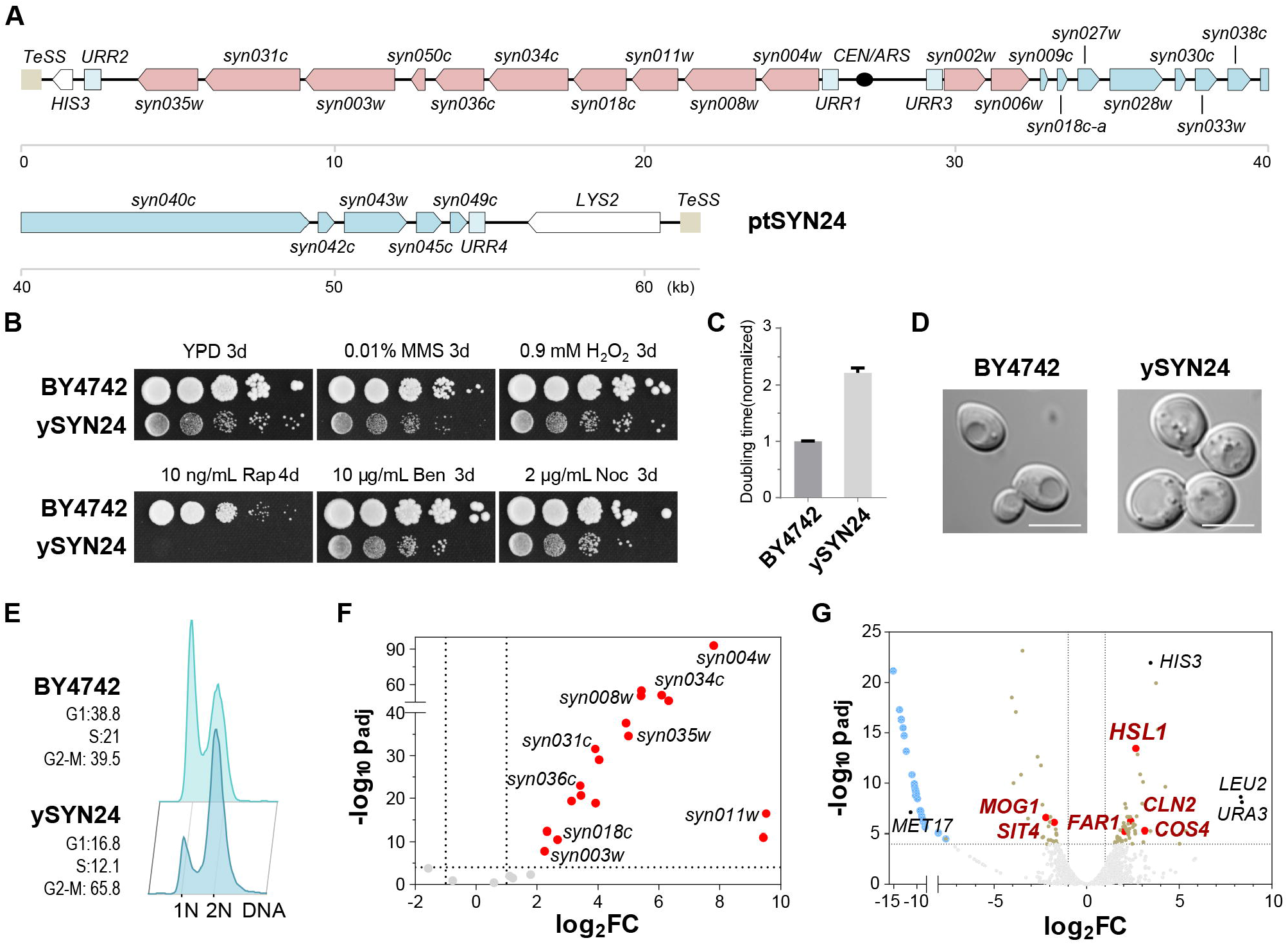
Functional replacement of *chrXIIL* with a completely revamped neochromosome. (A) Schematic representation of gene arrangement on ptSYN24 neochromosome. (B) Fitness of ySYN24 in different growth conditions. (C) The doubling time of ySYN24. (D) The cell morphology of ySYN24. Scale bars, 5 μm. (E) FACS analysis after propidine iodide staining on asynchronous cells. (F) Expression of genes on ptSYN24. Differentially expressed genes on ptSYN24 are marked in red. The nine TUs in yWT10 were labeled. (G) Transcript profiling of ySYN24. Genes in (F) were not shown in the volcano plot. Blue, genes deleted from *chrXIIL*; red, the six candidates responsible for the phenotypes of ySYN24; black, marker genes; brown, remaining differentially expressed genes.

Several characterizations were performed. At first, ySYN24 showed growth defects on YPD plates and severe sensitivity to rapamycin, while less dramatic fitness defects were detected under other conditions (Figure 5B). Consistently, its doubling time was more than 2-fold longer than that of wild type (Figure 5C). The cell size was slightly enlarged in ySYN24 (Figure 5D and Figure S5L). The fraction of cells with 2N DNA in log-phase population were significantly higher than that of BY4742 (Figure 5E), suggesting a cell-cycle defect. In addition, ySYN24 could not be arrested efficiently as wild type under nutrient depletion conditions (Figure S5M). Next, we determined the transcriptional profile of ySYN24 (Figure 5F-G). RNA-seq analysis showed that three quarters of genes on the neochromosome (18/24) were overexpressed (Figure 5F). Besides the genes on *chrXIIL* and markers, 82 additional genes were differentially expressed. Among them, overexpression of three genes (*HSL1, FAR1* and *CLN2*) and loss of function of *MOG1* were reported to increase cell size (Adames et al., 2015; Jorgensen et al., 2002; Sopko et al., 2006). Overexpression of *COS4* showed more than 3% shift in cells containing a 2N DNA content (Stevenson et al., 2001). *SIT4, HSL1* and the recoded *YLL003W* are involved in the G1/S or G2/M transition of mitotic cell cycle (Ma et al., 1999; McMillan et al., 1999; Sutton et al., 1991). Moreover, expression of these genes was not altered in yWT25 when compared to wild type (Table S3). Therefore, misexpression of these genes may collectively contribute to the cell cycle defect and abnormal cell size observed in ySYN24.

## DISCUSSION

The synthetic yeast genome has provided us a foundation to probe the function of eukaryotic genome with various design principles (Richardson et al., 2017). In this study, using *chrXIIL* as an example, we explored the possibility of simplifying the yeast genome by a combination of random deletions using SCRaMbLE and targeted knockout methods. We successfully removed over two-thirds of the sequences in the chromosome arm and generated a strain which could survive with only 12 genes (10 essential genes plus *YLL002W* and *YLL006W*). In addition, we demonstrated that each gene could be reconstructed using exogenous sequences without compromising its native function. In particular, we found that aggressive reprogramming of the coding sequences, that is, to encode each amino acid with the same codon, can be tolerated, at least on this small scale by the yeast. Together, these results suggest that the yeast genome is very plastic and tolerant to dramatic changes in gene composition, sequence and arrangement.

The successful synthesis of viral genomes, bacterial genomes and yeast chromosomes imply that chemically synthesized genomes are able to support life as well as the wild-type genomes (Cello et al., 2002; Fredens et al., 2019; Gibson et al., 2010; Luo et al., 2018b). In a further step, in this study, we demonstrated that the functions of wild-type chromosome can be assigned to neochromosomes with simplified gene contents, arrangements and sequences. The results in this study and several previous studies including relocating the rDNA locus (Mercy et al., 2017; Zhang et al., 2017) and minimizing chromosome number in yeast (Gu et al., 2022; Luo et al., 2018a; Shao et al., 2018) all suggest that the specific organizational form of yeast genome, including the chromosome number, gene organization and genome 3D structure, might only play minor, if there are any, roles in basic functions of yeast genome. This might be the consequence of a small genome and mainly short-range gene regulation in yeast (Jiang and Dai, 2019).

In nature, the 20 amino acids are encoded by 61 genetic codons, that is, most amino acids are encoded by more than one codon. Codon usage differs among organisms; and two of the 61 sense codons were removed from a synthesized *E. coli* genome (Fredens et al., 2019), suggesting the number of codons can be reduced. Theoretically, only 20 codons are needed to encode 20 amino acids. However, it may be difficult to achieve this minimal codon set, because of the potentially important roles of codon degeneracy in gene expression regulation (Komar, 2016). It is surprising that in this study most of the genes using the audacious rule that one amino acid is encoded by only one codon remain functional, suggesting the natural genome is of great plasticity.

To prevent unexpected deleterious effects on gene function and to reserve the wild-type like fitness in the synthetic strains, no sequence engineering was applied to the promoters and terminators in Sc2.0 (Richardson et al., 2017). Synthetic promoters and terminators designed based on the architecture of native ones yield diverse but reproducible expression levels of fluorescent proteins and exogenous enzymes, which helps a lot in deciphering the eukaryotic gene-regulatory logic and metabolic engineering (Curran et al., 2015; de Boer et al., 2020; Kotopka and Smolke, 2020; Liu et al., 2020; Redden and Alper, 2015). In this study, we showed that, although no sequence homology exists between these synthetic promoters and terminators and the native ones, they actually can also be used to build functional TUs, including the essential and nonessential ones, demonstrating the high plasticity of regulatory sequences.

In summary, as a pilot of the Sc3.0 project which aims to revamp and minimize the yeast genome (Dai et al., 2020), this study presents the most radical changes we have ever made to a chromosome arm. It indicates substantial plasticity of the yeast genome and suggests the feasibility of using computationally generated sequences to support viability. At the same time, the fitness defects of the strain with totally artificial sequences and the failure to reprogram several elements suggest that the strategies used in this study will be far too risky to scale up to whole-genome engineering. Due to the small amount of artificial regulatory elements available and limited knowledge of the complex co-regulation network, nature sequences from related organisms may represent a better long-term solution for the reprogramming of whole yeast genome.

## Supporting information

Table S1 The designed sequences of neochromosomes. Related to Figure 1, Figure 3 and Figure 5

Table S2 The information used to identify the nonessential gene set. Related to Figure 3

Table S3 Differentially expressed genes in yWT25. Related to Figure 3

Table S4 Sequences of reconstructed TUs. Related to Figure 4 and Figure S4.

Table S5 Oligonucleotides in this paper. Related to STAR Methods

Table S6. Strains in this paper. Related to STAR Methods.

## ACKNOWLEDGMENTS

We would like to thank Dr. Chuanle Xiao for help on the Iso-seq analysis. This work was supported by grants from National Key Research and Development Program of China (2018YFA0900100), National Natural Science Foundation of China (31725002, 31800069, 31800082 and 32122050), Shenzhen Science and Technology Program (KQTD20180413181837372), Guangdong Provincial Key Laboratory of Synthetic Genomics (2019B030301006), Guangdong Natural Science Funds for Distinguished Young Scholar (2021B1515020060), Bureau of International Cooperation, Chinese Academy of Sciences (172644KYSB20180022) and Shenzhen Outstanding Talents Training Fund. Work by J.D.B. was supported by US NSF grants MCB-1026068, MCB-1443299, MCB-1616111 and MCB-1921641. This work was also supported by UK Biotechnology and Biological Sciences Research Council (BBSRC) grants BB/M005690/1, BB/P02114X/1 and BB/W014483/1, and a Volkswagen Foundation “Life? Initiative” Grant (Ref. 94 771) to YC.

## AUTHOR CONTRIBUTIONS

JD conceived and designed the study. SJ, ZL, and JD supervised the study. SJ, ZL, KY, ZC, WY, HW, LC, ZZL, HG, LH, CZ, WZ, CHZ, YT, TL performed the experiments. SJ, ZL, KY, SZ, WY, MM, DS carried out the sequencing and analyzed the data. SJ, ZL, YM, YC, JDB and JD analyzed the data and wrote the manuscript. All authors contributed to review and editing of the manuscript.

## FIGURE LEGENDS

**Figure S1.**
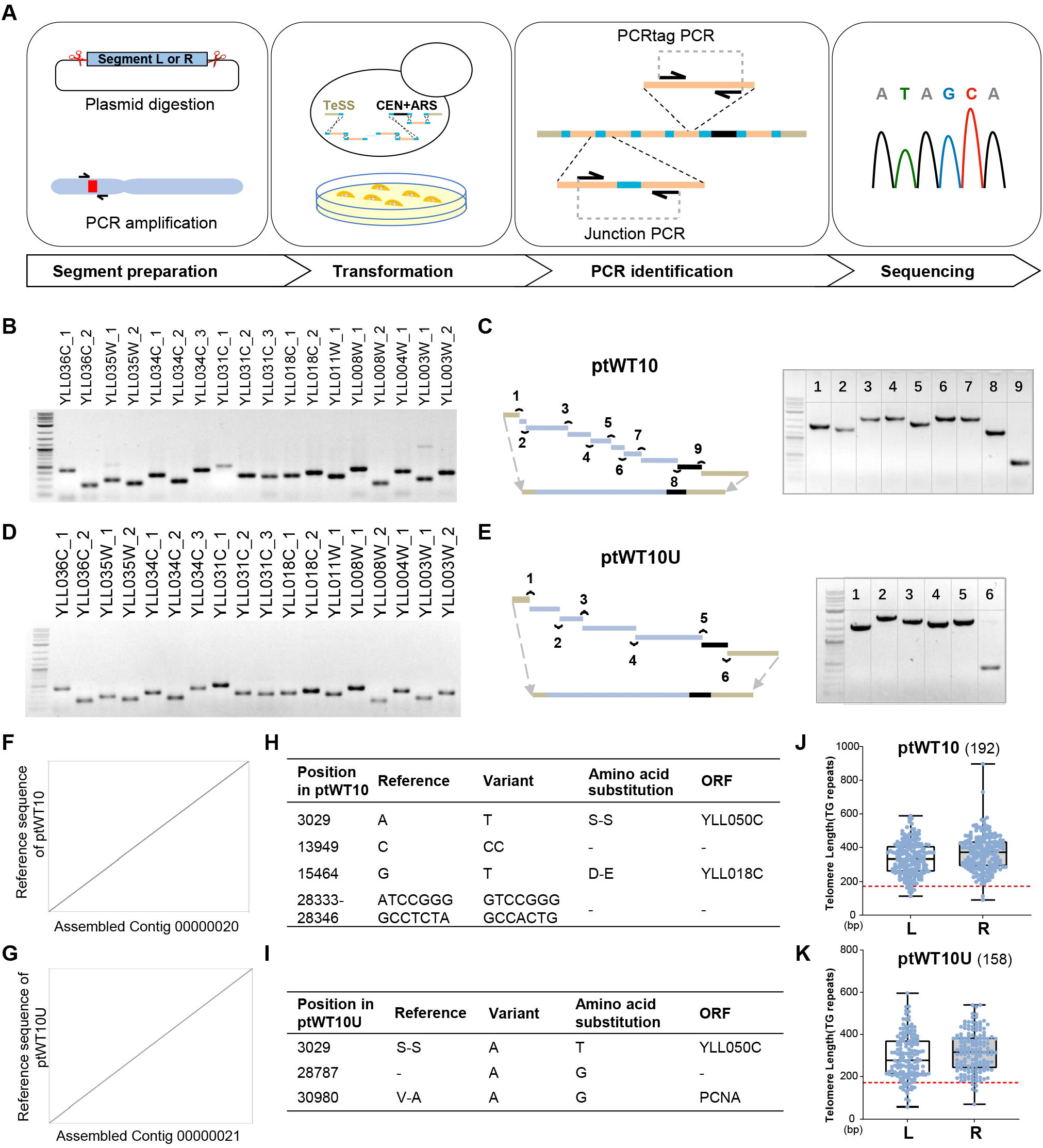
Construction of neochromosomes harboring essential genes from *chrXIIL*. Related to Figure 1. (A) The schematic workflow of transformation-assisted linear chromosome construction (TLCC). (B) Validation of ptWT10 neochromosome by PCRTag analysis. (C) Confirmation of assembled ptWT by junction analysis. The two ends and the black fragment represent segment L, R and M as in Fig 1A. (D) Validation of ptWT10U neochromosome. Similar analysis as in B. (E) Confirmation of ptWT10U. Similar analysis as C. (F/G) Dot plot analysis after whole genome sequencing showed the correct assembly of ptWT10 and ptWT10U. (H/I) Sequence variations of assembled ptWT10 and ptWT10U neochromosomes to the designed ones. (J-K) Length analysis of telomere TG repeats in neochromsomes. The red dashed line indicates the length of synthesized TG repeats. Only sequencing reads covering the entire neochromosomes (n=192 for ptWT10 and n=158 for ptWT10U) were used for calculation.

**Figure S2.**
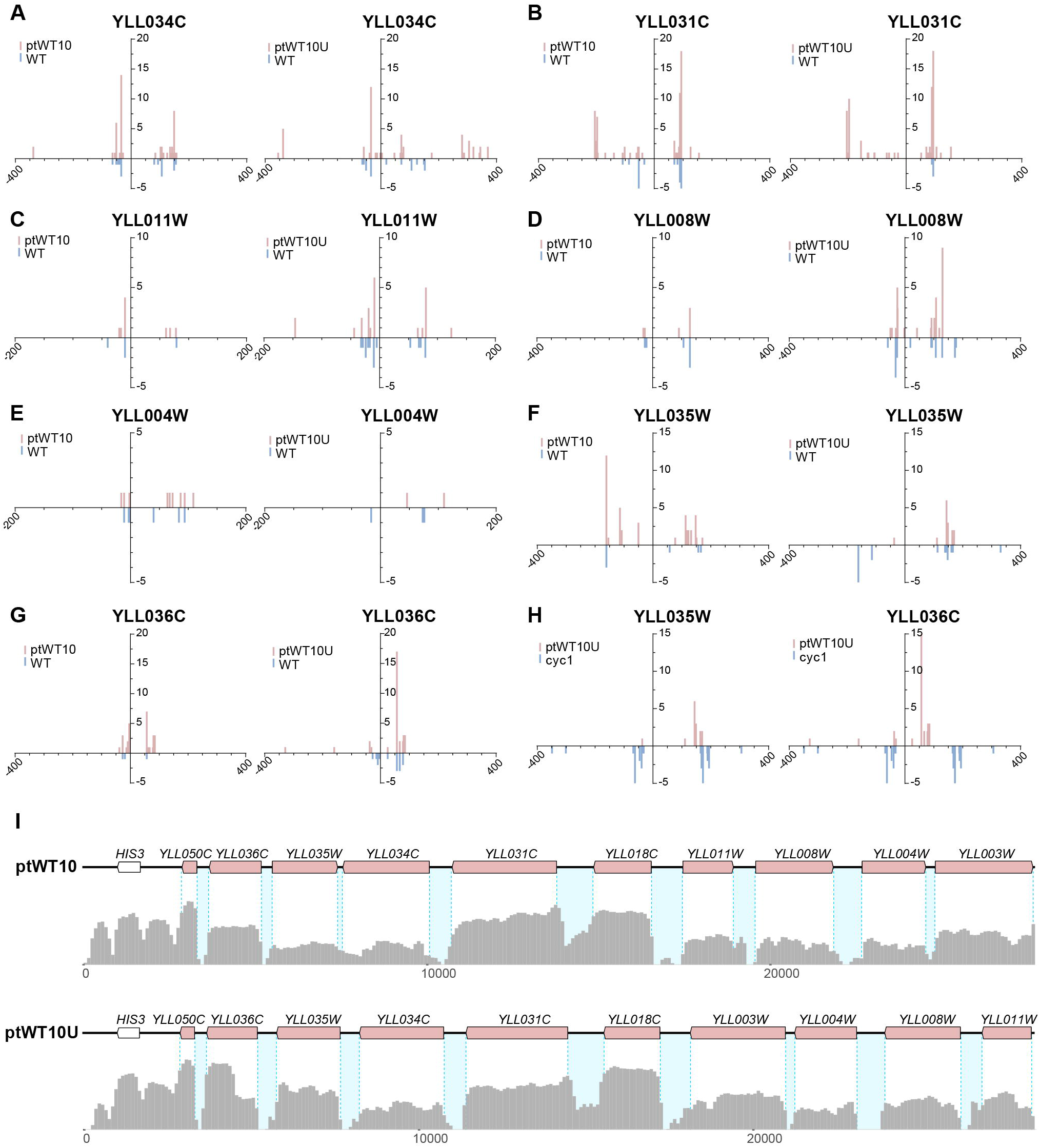
Transcription analysis of genes on ptWT10 and ptWT10U. Related to Figure 1. (A-B) The distribution of TSS and TES for the two genes, *YLL034C* and *YLL031C*, with the same direction on ptWT10 and ptWT10U. (C-E) The distribution of TSS and TES for other three genes, *YLL011W, YLL008W* and *YLL004W*, with different directions on ptWT10 and ptWT10U. (F-G) The distribution of TSS and TES for the two genes, *YLL035W* and *YLL036C*, with different regulation sequences on ptWT10 and ptWT10U. (H) Comparison of the distribution of TSS/TES of *YLL035W* and *YLL036C* on the ptWT10U to native *CYC1* gene. (I) Transcription expression of the essential genes on ptWT10 and ptWT10U in the yWT10 and yWT10U strains. Blue dashed lines, the boundaries of CDSs; light blue blocks, intergenic regions between essential genes. Y axis means log_10_ (average RNA sequencing depth of neo-chromosomes +1).

**Figure S3.**
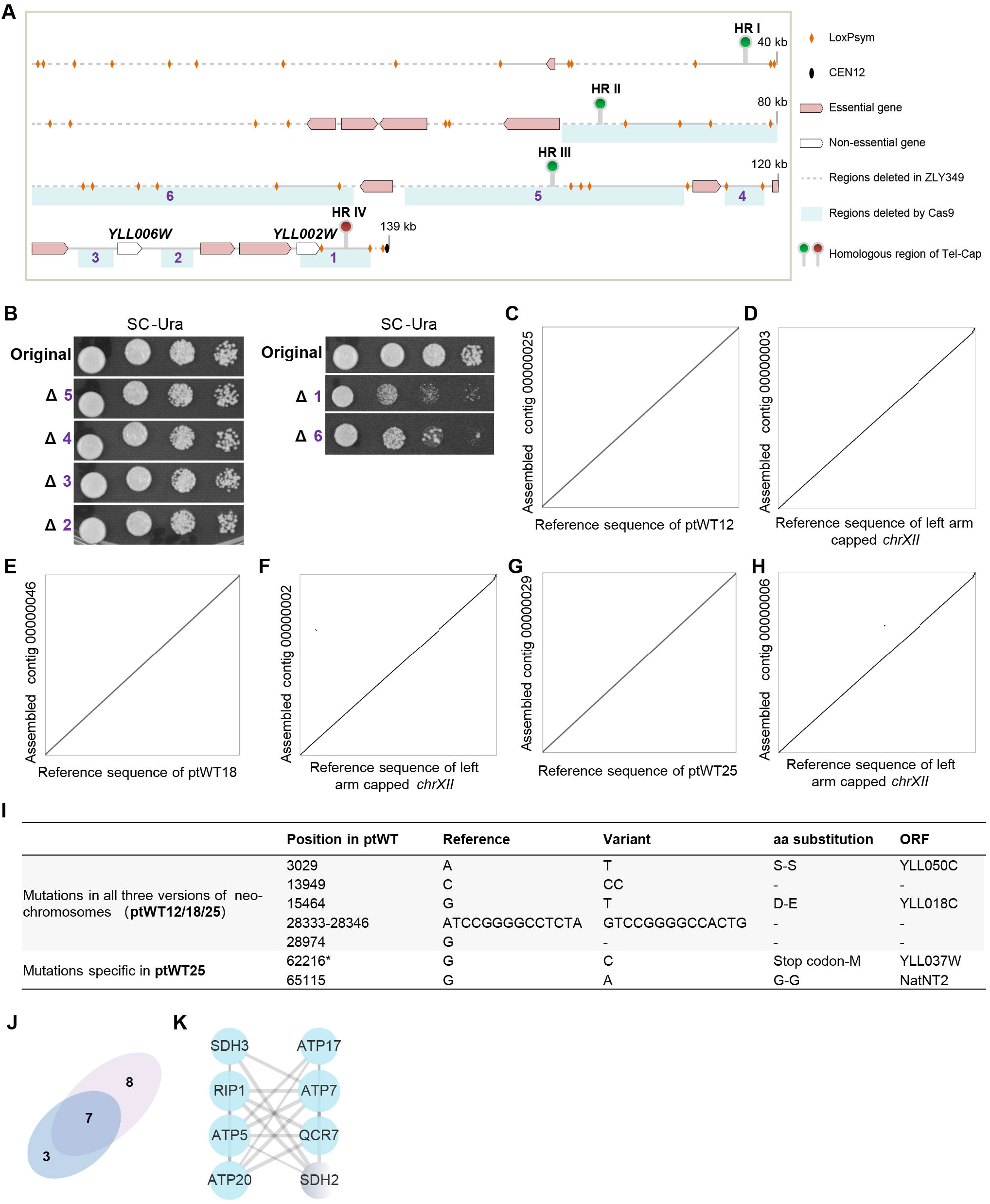
Simplification of *chrXIIL*. Related to Figure 2 and Figure 3. (A) The schematic diagram of left arm of *synXII*. Balloons (HR I-IV), positions of homologous regions used for Tel-Cap in Figure 2. (B) The growth of strains with corresponding regions deleted in (A). (C) Sequence comparison of neochromosome in yWT12 to the designed one. (D) Sequence analysis of *chrXII* in yWT12. (E) Sequence comparison of neochromosome in yWT18 to the designed one. (F) Sequence analysis of *chrXII* in yWT18. (G) Sequence comparison of neochromosome in yWT25 to the designed one. (H) Sequence analysis of *chrXII* in yWT25. (I) Sequence variations of assembled neochromosomes to the designed ones. (J) Venn diagram of the numbers of genes enriched in the oxidative phosphorylation pathway in Fig.3G and genes affecting cell growth when deleted which are down regulated in yWT25. (K) The interaction map between *SDH2* and the seven genes mentioned in (K). The network was generated with the string database (co-expression, cutoff score of 0.7). Node size, connection number.

**Figure S4.**
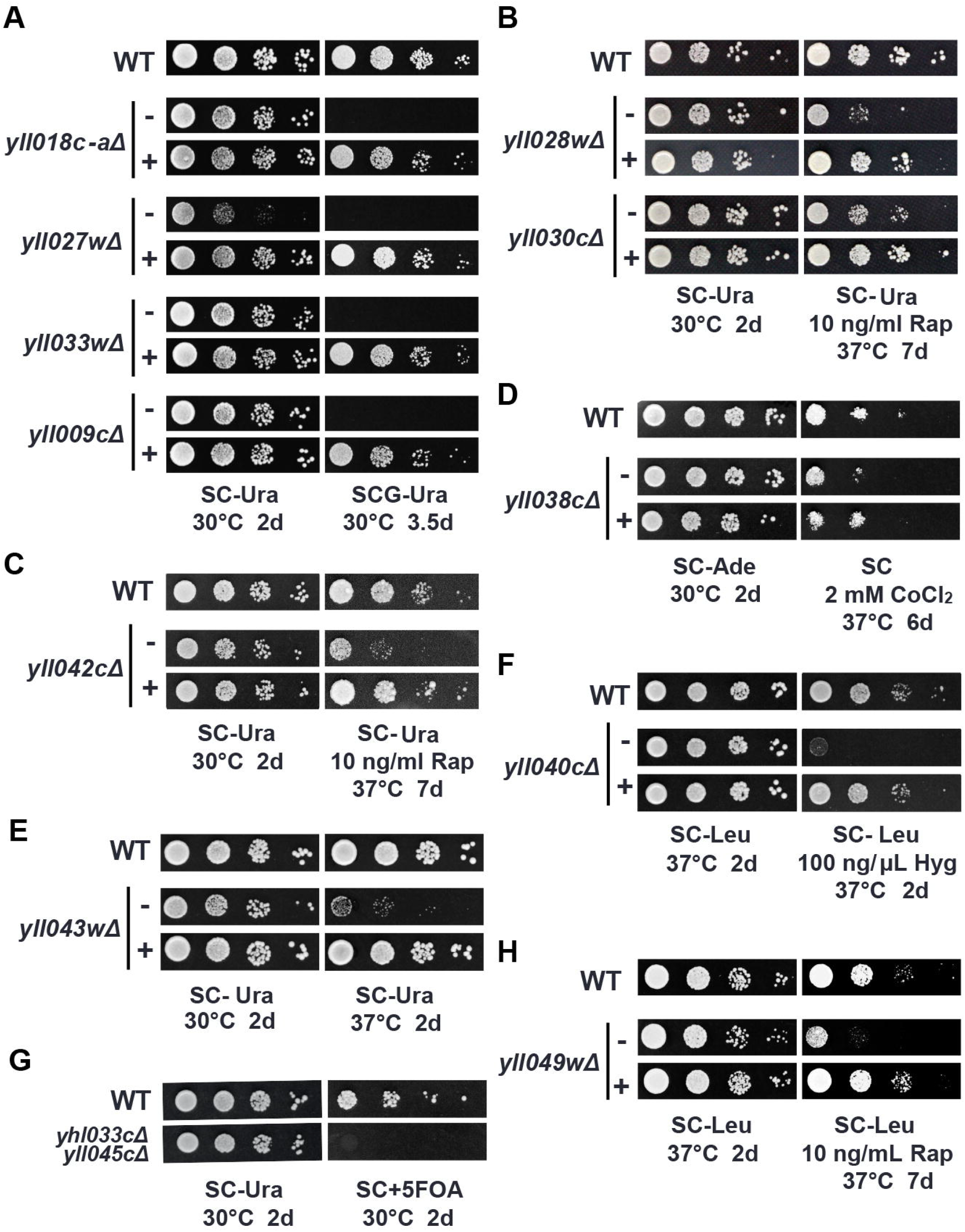
Reconstruct non-essential genes with artificial sequences. Related to Figure 4. (A-H) Serial dilution assay to measure the function of reconstructed nonessential genes. SCG, synthetic complete media with 3% glycerol. Hyg, hygromycin. In (D), *yll038c* deletion was reported to be sensitive to CoCl_2_ (Zhao et al., 2020). The limited sensitivity was rescued by the addition of reconstructed *syn038c*. In (G), *yhl033c* and *yll045c* double deletion is lethal (Steffen et al., 2012). -, without refactored gene; +, with refactored gene.

**Figure S5.**
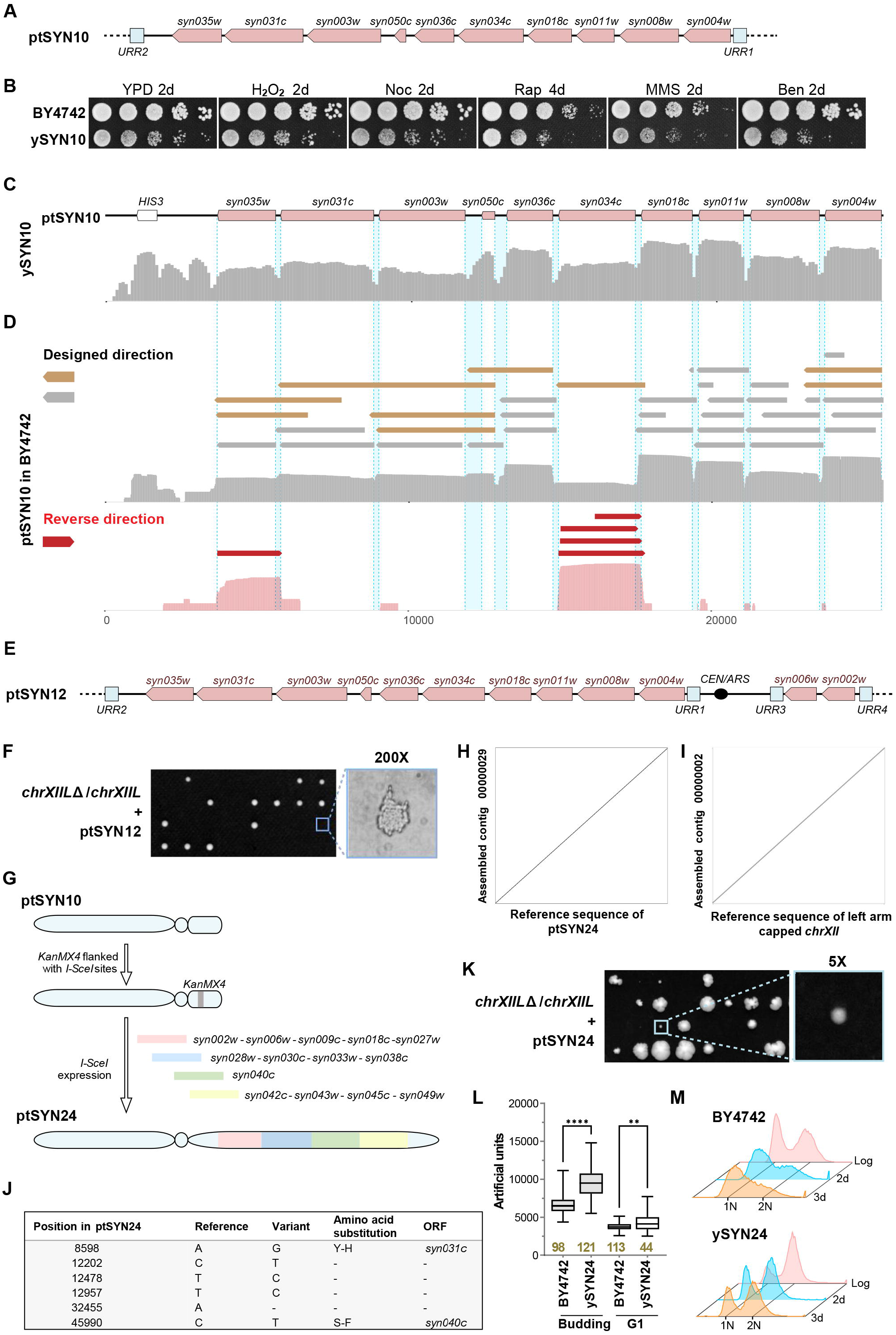
Construction of the neochromosomes with reconstructed genes. Related to Figure 5. (A)The schematic diagram of the reconstructed TUs added in ptSYN10. (B) Fitness analysis of the strain ySYN10. (C) Expression profile of genes from ptSYN10 in ySYN10. The RNA-seq data was shown in the same way as Figure S2I. (D) The Iso-seq results of the ptSYN10 in BY4742. The transcripts of each gene on ptSYN10, except *YLL050C*, were identified and divided into two categories: transcripts in designed direction and the ones in reverse direction. The identified isoforms for the synthetic genes were exhibited: the ones which went through more than two ORFs (brown), the anti-sense isoforms (red) and others (grey). (E) The schematic diagram of reconstructed TUs in ptSYN12. (F) Tetrad analysis of *chrXIIL*Δ/*chrXIIL* heterozygous strain containing ptSYN12. (G-J) Construction of ptSYN24. (H) Dot plot analysis suggested that the neochromosome was correctly assembled in *chrXIIL*Δ. (I) Dot plot analysis suggested that the left arm of *synXII* has been successfully deleted in these strains. (J) Sequence variations of assembled neochromosomes to the designed ones. (K) Tetrad analysis of c*hrXIIL*Δ/*chrXIIL* heterozygous strain with ptSYN24. (L) Quantification of cell size of ySYN24 strain. The box plot (min-max) shows cell size of budding cells and G1 cells (a single cell without bud). Cell sizes were measured by ImageJ. The number of cells counted for each strain were marked in brown. (M) FACS analysis of cells arrested by nutrient depletion. Cells in log phase culture were used as the control.

## STAR METHODS

### RESOURCE AVAILABILITY

#### Lead contact

Further information and request for reagents and resources should be directed to and will be fulfilled by the lead contact, Junbiao Dai.

#### Materials availability

All the requests for the generated plasmids and strains should be directed to the lead contact and will be made available on request after completion of a Materials Transfer Agreement.

#### Data and code availability

- Nanopore sequencing data, RNA-seq data and Iso-seq data have been deposited at SRA with bioproject number PRJNA883530 and are publicly available as of the date of publication.
- This paper does not report original code.
- Any additional information required to reanalyze the data reported in this paper is available from the lead contact upon request

#### EXPERIMENTAL MODEL AND SUBJECT DETAILS

##### Strains and growth media

The yeast strains used in this paper were derivates of BY4742 or *synXIIL(Luo et al*., *2021)*. Standard methods for yeast culture and transformation were applied. Targeted knockout methods were applied for deletion through homologous recombination. The strains generated in this study are listed in Table S6. For phenotypic analysis of strains, cells were cultured in YPD medium or YPD medium containing 0.9mM H_2_O_2_, 2μg/mL nocodazole, 10ng/mL rapamycin, 0.01% methyl methanesulfonate or 10μg/mL benomyl separately. For the function analysis of reconstructed TUs, cells were cultured on synthetic medium with or without additional drugs. Particularly, the carbon source in SCG-Ura was 3% glycerol, other than 2% glucose in other medium.

## METHOD DETAILS

### Construction of neochromosomes

The TLCC method includes four steps: segment preparation, yeast transformation, PCR identification and nanopore sequencing.

#### Segment preparation

All segments used for assembly contained at least 40 bp sequences overlapped with their adjacent segments at each side. The segments were released from plasmids using *NotI* digestion or PCR amplified using *synXIIL* genome as template.

(A) Totally 11 segments were used for assembling the ptWT10 neo-chromosome. Segment L and R were released from the constructed plasmids using *NotI* digestion, segment M was PCR amplified using a yeast strain containing a yeast artificial chromosome (Figure 1A). Segment 1-7 were amplified using synXIIL genome as template. The coordinates for these segments in the synXII reference sequence (Zhang et al., 2017) are: segment 1, encoding *YLL050C*, 27,316-28,330; segment 2, encoding *YLL034C, YLL035W* and *YLL036C*, 54,606-61,379; segment 3, encoding *YLL031C*, 64,787-68,394; segment 4, encoding *YLL018C*, 96,751-100,01; segment 5, encoding *YLL011W*, 115,173-117,239; segment 6, encoding *YLL008W*, 119,397-122,283; segment 7, encoding *YLL003W* and *YLL004W*, 128,529-134,272. In addition, a fused linker containing sequences of both URR3 and URR4 was also used to ligate segment M and R.

(B) Totally 8 segments were used for assembling the ptWT10U neo-chromosome. Segment L, R and M were prepared using the same method as ptWT10. Since the promoters of *YLL035W* and *YLL036C* share common sequence and some important elements of the promoter of *YLL036C* locate in the coding region of *YLL035W*, the promoters of *YLL035W* and *YLL036C* were changed to p*CYC1* and the terminator of *YLL035W* was changed to t*CYC1*. Segment 1-4 were amplified using the gDNA of BY4742 containing ptWT10 as template. The coordinates for these segments in the ptWT10 reference sequence are: segment 1, encoding *YLL050C* and *YLL036C*, 2,532-5,168; segment 2, encoding *YLL035W*, 5,492-7,390; segment 3, encoding *YLL034C, YLL031C* and *YLL018C*, 7,545-17,189; segment 4, encoding *YLL011W, YLL008W, YLL004W* and *YLL003W*, 17,190-27,887. The same fused linker in ptWT10 was also used.

(C) The ptWT12 was constructed by integrating a segment containing *YLL002W, YLL006W* and *KanMX4* at the right arm of ptWT10. The coordinates for *YLL002W* and *YLL006W* in *chrXII* (NC_001144.5) are: *YLL002W*, 146,041-147,802; *YLL006W*, 136,300-137,939.

(D) The ptWT18/25 were constructed similarly as ptWT12. The segment 1 containing 8 nonessential genes and a *KanMX4* selection marker was integrated between URR3 and URR4 of ptWT10 firstly (ptWT18), then the segment 2 containing 7 nonessential genes and a *NatNT2* selection marker was integrated at the middle of URR4 (ptWT25). The coordinates for these 15 nonessential genes in *chrXII* (NC_001144.5) are: *YLL002W*, 146,041-147,802; *YLL006W*, 136,300-137,939; *YLL009C*, 131,005-131,728; *YLL039W*, 63,593-65,707; *YLL040C*, 54,011-63,925; *YLL043W*, 49,438-52,087; *YLL045C*, 47,659-49,129; *YLL049W*, 40,179-41,381; *YLL018C-A*, 108,476-109,472; *YLL027W*, 86,903-88,356; *YLL028W*, 84,304-86,765; *YLL030C*, 80,156-81,197; *YLL033C*, 72,909-74,302; *YLL038C*, 65,575-67018; *YLL042C*, 51,877-53,090.

(E) Totally 7 segments are used for assembling the ptSYN10 neo-chromosome. Segment L, R and M are prepared using the same method as ptWT10 and ptWT10U. The segment 1-3 were released from the synthesized plasmids using *NotI* digestion. The same fused linker in ptWT10 was also used. The coordinates for these segments in the ptSYN10 reference sequence are: segment 1, encoding *syn035w* and *syn031c*, 2,536-8,907; segment 2, encoding *syn003w, syn050c, syn036c* and *syn034c*, 8,097-17,510; segment 3, encoding *syn018c, syn011w, syn008w* and *syn004w*,17,5111-25,503.

(F) Totally 7 segments were used for assembling the ptSYN12 neo-chromosome. Segment L, R and M were prepared using the same method as ptWT10 and ptWT10U. The segment 1-3 were prepared using the same method as ptSYN10. The segment 4 was the fused product, including URR3, URR4 and the DNA encoding *syn002w* and *syn006w*. The coordinates for these segments in the ptSYN12 reference sequence are: segment 1, encoding *syn035w* and *syn031c*, 2,536-8,907; segment 2, encoding *syn003w*, syn050c, *syn036c* and *syn034c*, 8,097-17,510; segment 3, encoding *syn018c, syn011w, syn008w* and *syn004w*, 17,511-25,503; segment 4, encoding *syn006w* and *syn002w*, 29,363-32,365.

(G) There were two steps to construct the ptSYN24 neo-chromosome based on ptSYN10. The first step integrated a segment containing *KanMX4* selection marker franked with two *I-SceI* sites between URR3 and URR4. Totally 6 segments for assembling the ptSYN24 neo-chromosome were transformed into the first-step strain expressing I-SceI restriction enzyme. Segment 1, 2, 4 were released from the plasmids containing synthetic genes by *BsaI* digestion. The segment 3 was amplified using the plasmid containing *syn040C*. Segment 5 and 6 were amplified by overlapping PCR. The coordinates for these segments in the ptSYN24 reference sequence are: segment 1, encoding *syn002w, syn006w, syn009c, syn018c* and *syn027w*, 28,813-34,513; Segment 2, encoding *syn028w, syn030c, syn033w* and *syn038c*, 34,474-39,327; Segment 3, encoding *syn040c*, 39,284-49,041; Segment 4, encoding *syn042c, syn043w, syn045c* and *syn049w*, 48,764-54,652; segment 5, 34,194-34,793; segment 6, 39,003-39,602.

#### Yeast transformation

the same protocol in our previous paper (Zhang et al., 2017) was used for yeast transformation. Briefly, a single colony was cultured in appropriate medium overnight and transferred to 5 mL appropriate medium with the OD_600_=0.1. The strains were harvested when the OD_600_ was between 0.4 and 0.6. Then the pellets were washed with ddH_2_O and 0.1 M LiOAc, respectively and 100μL 0.1 M LiOAc were used to resuspended the pellets and DNA fragments or plasmids were added. The mixture, including 312μL PEG3350, 41μL 1 M LiOAc and 25μL ssDNA (which were boiled at 100°C for 10min and cooled on the ice for 5min before used), were added and mixed. After standing 30min at 30°C incubator, 50μL DMSO were added and mix. Then the tube was subjected to the heat-shocker for 15min at 42°C. Cells were washed once with 5mM CaCl_2_ and plated onto the appropriate plates. The plates were cultured at 30°C for appropriate time.

#### PCR identification

primers for synthetic sequence identification (PCRtags) and junction verification were designed and listed in Table S5. The specificity of PCRtags were verified by using the genomic DNA of strains without corresponding neo-chromosome as template. PCR condition is the same as our previous report (Zhang et al., 2017). Briefly, in a 10μL reaction, 400ng genomic DNA were used. The reaction buffer and Easytaq® DNA polymerase were used according to the manufacturer’s instruction. Then carried out the following program in a thermal cycler: 1 cycle of 94°C for 5min, 30 cycles of 94°C for 30s, 55°C for 30s. 72°C for appropriate time, 1cycle of 72°C for 5min and 16°C keep. Only the colonies positive for all junctions and PCRtag were used for further sequencing.

#### Nanopore sequencing

The totally DNA was extracted using the QIAGEN Genomic-tips 100/G with Genomic DNA buffer Set using the manufacturer’s instruction. DNA quality was assessed by NanoDrop, gel-electrophoresis and Qubit. Libraries were prepared using the Ligation Sequencing Kit (SQK-LSK109) with the barcoding kits Native Barcoding Expansion 1-12 (PCR-free, EXP-NDB104) and Native Barcoding Expansion 13-24 (PCR-free, EXP-NDB104). Sequencing was performed using the MinlON platform with FLO-MIN106D for a 72-h running. Base calling, adaptor removal, and low-quality base filtering based on fast5 files were done by ONT software MinKNOW. The FastQ files were filtered by NanoFilt (De Coster et al., 2018) to remove short (read length < 500) and low-quality (average read quality score < 7) reads. The remaining reads were mapped to reference genome by NGMLR (Sedlazeck et al., 2018) and assembled into contigs using Canu 2.1.1 (Koren et al., 2017). Alignments between contigs and reference genomes were generated by MUMmer-nucmer 3.23 (Kurtz et al., 2004) with default parameters and were used to create collinearity dot plot by Dot (https://github.com/dnanexus/dot). The reads containing full-length neochromosome were extracted and aligned to reference sequence to infer the actual length of telomere TG repeats (TG repeats to the extremity).

### Pulsed-field gel electrophoresis and Southern blot

About 2 × 10^8^ cells at stationary phase were fixed in the 0.6% low melting point agarose for each plug and genomic DNA was prepared as described before (Zhang et al., 2017). Plug samples were resolved on a 1% agarose gel in 0.5 X TBE for 16 hours at 14°C on a BioRad CHEF Mapper XA Pulsed Field Electrophoresis System. The voltage was 6 V/cm, at an angle of 120° and switch time from initial 0.5 s to 1.5 s. The gels after PFGE were washed with ddH_2_O for twice and transferred onto Hybond-N+ membrane (Amersham) as described before. The samples were UV crosslinked and hybridized with DIG-labeled probes. Five DNA probes were amplified from corresponding linear chromosomes for probe labeling and mixed together for hybridization. Probe preparation, hybridization and detection were conducted with DIG-High Prime DNA Labeling and Detection Starter Kit (Roche, 11585614910). The primers for probes amplification were list in Table S5.

### Stability analysis of assembled chromosomes

Two independent colonies of the targeted strain were inoculated into 5 mL SC-His medium and cultured at 30°C with shaking at 220 rpm for 24 hrs. Then 5 μL of the culture was added into 5 mL fresh SC-His medium and grew for another 24 hrs. The cells after 10 days’ passages were plated onto SC-His plate to isolate single colonies. 10 colonies of each parent were picked and cultured separately for gDNA preparation and PCRtag analysis to see whether some deletion events have happened during about 125 generations.

Three out of the twenty colonies were further analyzed by whole genome sequencing using the Miseq platform. Because the neo-chromosomes are non-essential for BY4742, SC-His medium was used to select the neo-chromosome. The indicated strain was inoculated into 5 mL SC-His liquid medium and incubated at 30°C for 12 hours with shaking at 220 rpm. 5×10^7^ yeast cells were harvested for total DNA extraction using the Monarch® Genomic DNA Purification Kit based on the manufacturer’s instructions. 300-1000 bp libraries were prepared using the Nextera DNA Flex Library Prep Kit for purified genomic DNA and then sequenced on the Illumina Miseq platform with PE250 strategy. After adapter removal, the low-quality reads were trimmed by Trimmomatic (Bolger et al., 2014) with parameters “SLIDINGWINDOW:5:20 LEADING:5 TRAILING:5

MINLEN:50”, and the remaining reads were mapped to reference genome with BWA-mem (Li and Durbin, 2009). After marking PCR repeat using GATK-MarkDuplicates, the SNPs were called by GATK-HaplotypeCaller and GATK-GenotypeGVCFs. Structural variations were called using DELLY (Rausch et al., 2012).

### Serial dilution

The serial dilution was performed as previously mentioned (Zhang et al., 2017). In short, a single colony was inoculated into 3 mL YPD medium and incubated at 30°C with shaking at 220 rpm for 24 hours. The OD_600_ was measured, and the overnight cultures were diluted by sterile water to OD_600_=0.2. After four 10-fold gradient dilutions, well-mixed cells were dropped onto indicated plates. The plates were cultured at 30°C for appropriate time unless specifically mentioned.

### Transcriptome analysis

The non-stranded RNA sequencing libraries were prepared and sequenced by Beijing Novogene Bioinformatics Technology Co., Ltd. using Next® Ultra™ RNA Library Prep Kit for Illumina®. Because of the deletion of a few genes in our strains, we first generated the diminished reference genomes according to our design. We used Cutadapter software to remove adapters in raw data. HISAT2 and Picard were then used to accomplish the alignments of cleaned reads and remove PCR duplicates. Hereafter, Htseq-count software was employed to calculate read counts of each gene while intersection-nonempty option was set. The downstream statistical analysis was achieved by the DEseq2 package in R. Considering that the remaining fragment of *his3*Δ*1* in *chrXV* can still be transcribed, we only use the deleted part in *his3*Δ*1* to quantify *HIS3*. Differential expression in this paper was defined as |Log_2_ (Fold change) | > 1 and -log_10_ (Adjusted p-value) >4.

### Full-length transcriptome analysis

The transcriptomes of indicated strains were sequenced using the PacBio platform at Grandomics Company. Circular Consensus Sequencing (CCS) reads were generated using SMRT-Link (version 8.0.0.80529), with the following modified parameters: “--min-passes 0 --min-length 50 --max-length 21000 --min-rq 0.75”. We used Lima (version 1.10.0, commit SL-release-8.0.0) for Single Cell Full-Length Non-Concatemer (FLNC) reads detection, it’s integrated into the PacBio official SMRT-Link (version 8.0.0.80529) software package. Lima map 5’ and 3’ primers to CCS reads first, then parse standard pair of 5’ and 3’ primers CCS as the full-length isoform, next trim the primer sequence and polyA tail in each full-length isoform. Here, each isoform was oriented and correspond to cDNA orientation from 5’ to 3’ end. After FLNC detection, primer and polyA tail trimming, the remaining fraction of each isoform FLNC read was aligned to the reference genome with minimap2 (version 2.17-r974-dirty) in spliced alignment mode with parameters: “-ax splice -uf --secondary=no - C5”. Reads with more than 95% identity and 30% coverage of a certain CDS are directly counted from paf files.

To ensure the generation of transcripts with high accuracy, we use cDNA_Cupcake (https://github.com/Magdoll/cDNA_Cupcake) python script “collapse_isoforms_by_sam.py” to collapse redundant isoforms. The “--flnc-coverage” for minimum collapsed reads is set to 5 and the other parameters are set to default.

After redundant isoforms collapsing, unique isoforms can be reported as GFF file. A homemade R script is used to illustrate the TSS, TES and transcription direction based on gggenes (https://github.com/wilkox/gggenes).

### Tel-cap

The four homologous fragments were amplified from the *synXIIL* and respectively cloned into the plasmid with TeSS sequences using the restriction sites, *XmaI* and *SalI*. The TeSS-Marker-HR fragments were released by *BsaI* digestion and transformed into the heterozygous diploid cells (*chrXII* X synXIIL). The chromosome arm capped strains were screened with the PCRtags which have been reported to distinguish *chrXIIL*(1.0) and *synXIIL*(2.0) (Zhang et al., 2017). The colonies which were positive for 1.0 PCRtags but negative for 2.0 PCRtags indicated the specific loss of sequences on *synXIIL*.

### Tetrad analysis

Diploid strains were cultured in selective medium at 30°C overnight. About 8 × 10^7^ cells were harvested and washed with ddH_2_O twice. Cells were resuspended with 50 μL 1 × Sporulation medium (10 g/L potassium acetate, 0.05 g/L zinc acetate dehydrate) and transferred into 2 ml 1 × Sporulation medium. The tubes were incubated at 25°C for 3-10 days. Then cells were harvested and resuspended in 30 μL Zymolyase-100T (0.5 mg/ml Zymolyase-100T in 1 M sorbitol) for about 4-6 minutes at RT. Add 300 μL pre-cold ddH_2_O to stop digestion. Spread 20 μL suspension on YPD plates gently for further dissection under the microscope. These plates were cultured at 30°C for appropriate days before imaging, replicated onto various selective media to identify their auxotroph and mating type.

### Interaction analysis

All genetic interactions of chromosome XII-left arm genes of *Saccharomyces cerevisiae* are available at the Saccharomyces Genome Database (SGD) (https://www.yeastgenome.org). In this study, the genetic interactions were counted until 23 April 2021, and self-interactions of a gene were eliminated before calculation in Table S2.

As for the network analysis for the differentially expressed genes in Figure 3 and S3K, the interaction map was generated using the string database with different parameters (Szklarczyk et al., 2021) and visualized by Cytoscape 3.7.2.

### YeastFab assembly and screening of functional non-essential TUs

The pMV-*AmpR* plasmids, ORF plasmids and HCKan_terminator plasmids, hosting promoter pool (P), ORF (O) and terminator pool (T) were prepared for YeastFab assembly. The promoter fragments were amplified by ExTaq DNA polymerase (TaKaRa). The YeastFab assembly is performed according to the reaction system below: 1 μL 10X Buffer for T4 DNA Ligase (NEB), 0.1 μL Purified BSA 100X (Thermo), 0.2 μL T4 DNA Ligase (Thermo), 0.5 μL Esp3 I (Thermo), 2 μL purified promoter PCR product, 2 μL ORF plasmid (20 ng/μL), 2 μL HCKan_terminator plasmid (20 ng/μL), 2 μL vector plasmid (20 ng/μL) and 0.2 μL ddH_2_O. Then carry out the following program in a thermal cycler: 37°C for 2 h, 55°C for 15 min, 80°C for 15 min. The reaction products were transformed into *E. coli* DH5α and 5 mL LB liquid medium containing carbenicillin disodium (100 μg/mL) was added into the bacteria mix, shaking at 37°C overnight. Plasmid pool was extracted from the bacteria mixture and transformed into yeast. Then the yeast cells were plated on selective plates.

Six single colonies were randomly picked from the plates and cultured in selective liquid media overnight at 30°C. The OD_600_ was measured, and the overnight cultures were diluted by sterile water to OD_600_=1. After four 10-fold gradient dilution, well-mixed cells were dropped onto indicated plates. The plates were cultured at appropriate temperature. The yeast strains with recovered phenotypes were selected. The plasmids carrying the synthetic TUs were isolated and identified by sanger sequencing.

### Flow cytometry analysis

Samples were selected at corresponding time points and fixed cells with 70% ethanol overnight at 4°C. Cells were resuspended in 50mM sodium citrate (pH 7.0) and sonicated on ice shortly. Cells were resuspended in 50 mM sodium citrate (pH 7.0) and added with RnaseA (0.25 mg/mL) for incubation at 37°C. Cells were washed with 50mM sodium citrate (pH 7.0) and resuspended in 50mM sodium citrate (pH 7.0) containing propidium iodide (16 µg/mL). The cells were incubated at room temperature for at least 1 hour. Samples were proceeded with BD FACS Celesta for measurement. The software FlowJo was used for analysis and the fraction of cells at different stages was calculated with Dean-Jett-Fox model.

### Microscope imaging

Cells were cultured in YPD overnight and subcultured into fresh YPD for several hours to get cells at log phase. Cells were harvested gently and washed once with ddH_2_O. Then cells were dropped on slides for further imaging with a Nikon A1 confocal microscope under 60 × objective. To calculate cell size of strains at different stages with Image J, the cells of each strain were defined into two groups: G1 cells (not budding) and dividing cells (budding).

## QUANTIFICATION AND STATISTICAL ANALYSIS

Error bars represent SD. The data was analyzed using MS-Excel. Two-tailed t-tests were used to compare different groups in this paper. Differences were considered as statistically significant at p-value<0.05. * P<0.05, ** P<0.01, *** P<0.001, **** P<0.0001.

**Table S1. The designed sequences of neochromosomes. Related to Figure 1**,**3 and 5**

**Table S2. The information used to identify the nonessential gene set. Related to Figure 3**.

**Table S3. Differentially expressed genes in yWT25. Related to Figure 3**.

**Table S4. Sequences of reconstructed TUs. Related to Figure 4 and S4**.

**Table S5. Oligonucleotides in this paper. Related to STAR Methods**.

**Table S6. Strains in this paper. Related to STAR Methods**.

## REFERENCE

Adames, N.R., Schuck, P.L., Chen, K.C., Murali, T.M., Tyson, J.J., and Peccoud, J. (2015). Experimental testing of a new integrated model of the budding yeast Start transition. Mol Biol Cell 26, 3966–3984.

Annaluru, N., Muller, H., Mitchell, L.A., Ramalingam, S., Stracquadanio, G., Richardson, S.M., Dymond, J.S., Kuang, Z., Scheifele, L.Z., Cooper, E.M., et al. (2014). Total synthesis of a functional designer eukaryotic chromosome. Science 344, 55–58.

Blattner, F.R., Plunkett, G., 3rd, Bloch, C.A., Perna, N.T., Burland, V., Riley, M., Collado-Vides, J., Glasner, J.D., Rode, C.K., Mayhew, G.F., et al. (1997). The complete genome sequence of Escherichia coli K-12. Science 277, 1453–1462.

Bolger, A.M., Lohse, M., and Usadel, B. (2014). Trimmomatic: a flexible trimmer for Illumina sequence data. Bioinformatics 30, 2114–2120.

Brooks, A.N., Hughes, A.L., Clauder-Munster, S., Mitchell, L.A., Boeke, J.D., and Steinmetz, L.M. (2022). Transcriptional neighborhoods regulate transcript isoform lengths and expression levels. Science 375, 1000–1005.

Cello, J., Paul, A.V., and Wimmer, E. (2002). Chemical synthesis of poliovirus cDNA: generation of infectious virus in the absence of natural template. Science 297, 1016–1018.

Curran, K.A., Morse, N.J., Markham, K.A., Wagman, A.M., Gupta, A., and Alper, H.S. (2015). Short Synthetic Terminators for Improved Heterologous Gene Expression in Yeast. ACS Synth Biol 4, 824–832.

Dai, J., Boeke, J.D., Luo, Z., Jiang, S., and Cai, Y. (2020). Sc3.0: revamping and minimizing the yeast genome. Genome Biology 21, 205.

Dani, G.M., and Zakian, V.A. (1983). Mitotic and meiotic stability of linear plasmids in yeast. Proc Natl Acad Sci U S A 80, 3406–3410.

de Boer, C.G., Vaishnav, E.D., Sadeh, R., Abeyta, E.L., Friedman, N., and Regev, A. (2020). Deciphering eukaryotic gene-regulatory logic with 100 million random promoters. Nat Biotechnol 38, 56–65.

De Coster, W., D’Hert, S., Schultz, D.T., Cruts, M., and Van Broeckhoven, C. (2018). NanoPack: visualizing and processing long-read sequencing data. Bioinformatics 34, 2666–2669.

Driscoll, R., Hudson, A., and Jackson, S.P. (2007). Yeast Rtt109 promotes genome stability by acetylating histone H3 on lysine 56. Science 315, 649–652.

Dymond, J.S., Richardson, S.M., Coombes, C.E., Babatz, T., Muller, H., Annaluru, N., Blake, W.J., Schwerzmann, J.W., Dai, J.B., Lindstrom, D.L., et al. (2011). Synthetic chromosome arms function in yeast and generate phenotypic diversity by design. Nature 477, 471–U124.

Finley, D., Ozkaynak, E., and Varshavsky, A. (1987). The Yeast Polyubiquitin Gene Is Essential for Resistance to High-Temperatures, Starvation, and Other Stresses. Cell 48, 1035–1046.

Fourel, G., Revardel, E., Koering, C.E., and Gilson, E. (1999). Cohabitation of insulators and silencing elements in yeast subtelomeric regions. Embo j 18, 2522–2537.

Fraser, C.M., Gocayne, J.D., White, O., Adams, M.D., Clayton, R.A., Fleischmann, R.D., Bult, C.J., Kerlavage, A.R., Sutton, G., Kelley, J.M., et al. (1995). The minimal gene complement of Mycoplasma genitalium. Science 270, 397–403.

Fredens, J., Wang, K., de la Torre, D., Funke, L.F.H., Robertson, W.E., Christova, Y., Chia, T., Schmied, W.H., Dunkelmann, D.L., Beranek, V., et al. (2019). Total synthesis of Escherichia coli with a recoded genome. Nature 569, 514–518.

Gibson, D.G., Glass, J.I., Lartigue, C., Noskov, V.N., Chuang, R.Y., Algire, M.A., Benders, G.A., Montague, M.G., Ma, L., Moodie, M.M., et al. (2010). Creation of a bacterial cell controlled by a chemically synthesized genome. Science 329, 52–56.

Glass, J.I., Assad-Garcia, N., Alperovich, N., Yooseph, S., Lewis, M.R., Maruf, M., Hutchison, C.A., 3rd, Smith, H.O., and Venter, J.C. (2006). Essential genes of a minimal bacterium. Proc Natl Acad Sci U S A 103, 425–430.

Goffeau, A., Barrell, B.G., Bussey, H., Davis, R.W., Dujon, B., Feldmann, H., Galibert, F., Hoheisel, J.D., Jacq, C., Johnston, M., et al. (1996). Life with 6000 genes. Science 274, 546, 563-547.

Gu, X., Ye, T., Zhang, X.R., Nie, L., Wang, H., Li, W., Lu, R., Fu, C., Du, L.L., and Zhou, J.Q. (2022). Single-chromosome fission yeast models reveal the configuration robustness of a functional genome. Cell Rep 40, 111237.

Guo, Y., Dong, J., Zhou, T., Auxillos, J., Li, T., Zhang, W., Wang, L., Shen, Y., Luo, Y., Zheng, Y., et al. (2015). YeastFab: the design and construction of standard biological parts for metabolic engineering in Saccharomyces cerevisiae. Nucleic Acids Res 43, e88.

Han, J., Zhou, H., Horazdovsky, B., Zhang, K., Xu, R.-M., and Zhang, Z. (2007). Rtt109 Acetylates Histone H3 Lysine 56 and Functions in DNA Replication. Science 315, 653–655.

Hanway, D., Chin, J.K., Xia, G., Oshiro, G., Winzeler, E.A., and Romesberg, F.E. (2002). Previously uncharacterized genes in the UV-and MMS-induced DNA damage response in yeast. Proc Natl Acad Sci U S A 99, 10605–10610.

Hutchison, C.A., 3rd, Chuang, R.Y., Noskov, V.N., Assad-Garcia, N., Deerinck, T.J., Ellisman, M.H., Gill, J., Kannan, K., Karas, B.J., Ma, L., et al. (2016). Design and synthesis of a minimal bacterial genome. Science 351, aad6253.

Isaacs, F.J., Carr, P.A., Wang, H.H., Lajoie, M.J., Sterling, B., Kraal, L., Tolonen, A.C., Gianoulis, T.A., Goodman, D.B., Reppas, N.B., et al. (2011). Precise manipulation of chromosomes in vivo enables genome-wide codon replacement. Science 333, 348–353.

Jensen, L.T., and Culotta, V.C. (2000). Role of Saccharomyces cerevisiae ISA1 and ISA2 in iron homeostasis. Mol Cell Biol 20, 3918–3927.

Jiang, S., Zhao, S., Cai, Z., Tang, Y., and Dai, J. (2020a). Synthetic yeast genomes for studying chromosomal features. Current Opinion in Systems Biology 23, 1–7.

Jiang, S.Y., and Dai, J.B. (2019). Inevitability or contingency: how many chromosomes do we really need? Science China-Life Sciences 62, 140–143.

Jiang, S.Y., Si, T., and Dai, J.B. (2020b). Whole-Genome Regulation for Yeast Metabolic Engineering. Small Methods 4.

Jorgensen, P., Nishikawa, J.L., Breitkreutz, B.J., and Tyers, M. (2002). Systematic identification of pathways that couple cell growth and division in yeast. Science 297, 395–400.

Koike, N., Hatano, Y., and Ushimaru, T. (2018). Heat shock transcriptional factor mediates mitochondrial unfolded protein response. Curr Genet 64, 907–917.

Komar, A.A. (2016). The Yin and Yang of codon usage. Hum Mol Genet 25, R77–r85.

Koren, S., Walenz, B.P., Berlin, K., Miller, J.R., Bergman, N.H., and Phillippy, A.M. (2017). Canu: scalable and accurate long-read assembly via adaptive k-mer weighting and repeat separation. Genome research 27, 722–736.

Kornmann, B., Currie, E., Collins, S.R., Schuldiner, M., Nunnari, J., Weissman, J.S., and Walter, P. (2009). An ER-mitochondria tethering complex revealed by a synthetic biology screen. Science 325, 477–481.

Kotopka, B.J., and Smolke, C.D. (2020). Model-driven generation of artificial yeast promoters. Nat Commun 11, 2113.

Kurtz, S., Phillippy, A., Delcher, A.L., Smoot, M., Shumway, M., Antonescu, C., and Salzberg, S.L. (2004). Versatile and open software for comparing large genomes. Genome biology 5, R12.

Lajoie, M.J., Rovner, A.J., Goodman, D.B., Aerni, H.R., Haimovich, A.D., Kuznetsov, G., Mercer, J.A., Wang, H.H., Carr, P.A., Mosberg, J.A., et al. (2013). Genomically recoded organisms expand biological functions. Science 342, 357–360.

Li, H., and Durbin, R. (2009). Fast and accurate short read alignment with Burrows-Wheeler transform. Bioinformatics 25, 1754–1760.

Liang, Z., Luo, Z., Zhang, W., Yu, K., Wang, H., Geng, B., Yang, Q., Ni, Z., Zeng, C., Zheng, Y., et al. (2022). Synthetic refactor of essential genes decodes functionally constrained sequences in yeast genome. iScience 25, 104982.

Liu, R., Liu, L., Li, X., Liu, D., and Yuan, Y. (2020). Engineering yeast artificial core promoter with designated base motifs. Microb Cell Fact 19, 38.

Luo, J., Sun, X., Cormack, B.P., and Boeke, J.D. (2018a). Karyotype engineering by chromosome fusion leads to reproductive isolation in yeast. Nature 560, 392–396.

Luo, Z., Yang, Q., Geng, B., Jiang, S., Yang, S., Li, X., Cai, Y., and Dai, J. (2018b). Whole genome engineering by synthesis. Sci China Life Sci 61, 1515–1527.

Luo, Z., Yu, K., Xie, S., Monti, M., Schindler, D., Fang, Y., Zhao, S., Liang, Z., Jiang, S., Luan, M., et al. (2021). Compacting a synthetic yeast chromosome arm. Genome Biol 22, 5.

Ma, P.S., Winderickx, J., Nauwelaers, D., Dumortier, F., De Doncker, A., Thevelein, J.M., and Van Dijck, P. (1999). Deletion of SFI1, a novel suppressor of partial Ras-cAMP pathway deficiency in the yeast Saccharomyces cerevisiae, causes G(2) arrest. Yeast 15, 1097–1109.

McMillan, J.N., Longtine, M.S., Sia, R.A.L., Theesfeld, C.L., Bardes, E.S.G., Pringle, J.R., and Lew, D.J. (1999). The morphogenesis checkpoint in Saccharomyces cerevisiae: Cell cycle control of Swe1p degradation by Hsl1p and Hsl7p. Molecular and Cellular Biology 19, 6929–6939.

Mercy, G., Mozziconacci, J., Scolari, V.F., Yang, K., Zhao, G., Thierry, A., Luo, Y., Mitchell, L.A., Shen, M., Shen, Y., et al. (2017). 3D organization of synthetic and scrambled chromosomes. Science 355.

Miller, M.A., Russo, J., Fischer, A.D., Leban, F.A.L., and Olivas, W.M. (2014). Carbon source-dependent alteration of Puf3p activity mediates rapid changes in the stabilities of mRNAs involved in mitochondrial function. Nucleic Acids Research 42, 3954–3970.

Murakami, K., Tao, E., Ito, Y., Sugiyama, M., Kaneko, Y., Harashima, S., Sumiya, T., Nakamura, A., and Nishizawa, M. (2007). Large scale deletions in the Saccharomyces cerevisiae genome create strains with altered regulation of carbon metabolism. Applied Microbiology and Biotechnology 75, 589–597.

Murray, A.W., and Szostak, J.W. (1983). Construction of artificial chromosomes in yeast. Nature 305, 189–193.

Olivas, W., and Parker, R. (2000). The Puf3 protein is a transcript-specific regulator of mRNA degradation in yeast. Embo Journal 19, 6602–6611.

Pelletier, J.F., Sun, L., Wise, K.S., Assad-Garcia, N., Karas, B.J., Deerinck, T.J., Ellisman, M.H., Mershin, A., Gershenfeld, N., Chuang, R.Y., et al. (2021). Genetic requirements for cell division in a genomically minimal cell. Cell 184, 2430–2440 e2416.

Posfai, G., Plunkett, G., 3rd, Feher, T., Frisch, D., Keil, G.M., Umenhoffer, K., Kolisnychenko, V., Stahl, B., Sharma, S.S., de Arruda, M., et al. (2006). Emergent properties of reduced-genome Escherichia coli. Science 312, 1044–1046.

Rausch, T., Zichner, T., Schlattl, A., Stutz, A.M., Benes, V., and Korbel, J.O. (2012). DELLY: structural variant discovery by integrated paired-end and split-read analysis. Bioinformatics 28, i333–i339.

Ray, A., and Runge, K.W. (1999). The yeast telomere length counting machinery is sensitive to sequences at the telomere-nontelomere junction. Molecular and Cellular Biology 19, 31–45.

Redden, H., and Alper, H.S. (2015). The development and characterization of synthetic minimal yeast promoters. Nature Communications 6, 7810.

Richardson, S.M., Mitchell, L.A., Stracquadanio, G., Yang, K., Dymond, J.S., DiCarlo, J.E., Lee, D., Huang, C.L., Chandrasegaran, S., Cai, Y., et al. (2017). Design of a synthetic yeast genome. Science 355, 1040–1044.

Richardson, S.M., Nunley, P.W., Yarrington, R.M., Boeke, J.D., and Bader, J.S. (2010). GeneDesign 3.0 is an updated synthetic biology toolkit. Nucleic Acids Research 38, 2603–2606.

Sedlazeck, F.J., Rescheneder, P., Smolka, M., Fang, H., Nattestad, M., von Haeseler, A., and Schatz, M.C. (2018). Accurate detection of complex structural variations using single-molecule sequencing. Nat Methods 15, 461-+.

Shao, Y., Lu, N., Wu, Z., Cai, C., Wang, S., Zhang, L.L., Zhou, F., Xiao, S., Liu, L., Zeng, X., et al. (2018). Creating a functional single-chromosome yeast. Nature 560, 331–335.

Sopko, R., Huang, D., Preston, N., Chua, G., Papp, B., Kafadar, K., Snyder, M., Oliver, S.G., Cyert, M., Hughes, T.R., et al. (2006). Mapping pathways and phenotypes by systematic gene overexpression. Mol Cell 21, 319–330.

Steffen, K.K., McCormick, M.A., Pham, K.M., MacKay, V.L., Delaney, J.R., Murakami, C.J., Kaeberlein, M., and Kennedy, B.K. (2012). Ribosome deficiency protects against ER stress in Saccharomyces cerevisiae. Genetics 191, 107–118.

Stevenson, L.F., Kennedy, B.K., and Harlow, E. (2001). A large-scale overexpression screen in Saccharomyces cerevisiae identifies previously uncharacterized cell cycle genes. Proceedings of the National Academy of Sciences of the United States of America 98, 3946–3951.

Sutton, A., Immanuel, D., and Arndt, K.T. (1991). The Sit4 Protein Phosphatase Functions in Late G1 for Progression into S-Phase. Molecular and Cellular Biology 11, 2133–2148.

Szklarczyk, D., Gable, A.L., Nastou, K.C., Lyon, D., Kirsch, R., Pyysalo, S., Doncheva, N.T., Legeay, M., Fang, T., Bork, P., et al. (2021). The STRING database in 2021: customizable protein-protein networks, and functional characterization of user-uploaded gene/measurement sets. Nucleic Acids Res 49, D605–D612.

Yoshikawa, K., Tanaka, T., Ida, Y., Furusawa, C., Hirasawa, T., and Shimizu, H. (2011). Comprehensive phenotypic analysis of single-gene deletion and overexpression strains of Saccharomyces cerevisiae. Yeast 28, 349–361.

Zhang, W., Zhao, G., Luo, Z., Lin, Y., Wang, L., Guo, Y., Wang, A., Jiang, S., Jiang, Q., Gong, J., et al. (2017). Engineering the ribosomal DNA in a megabase synthetic chromosome. Science 355.

Zhao, Y.Y., Cao, C.L., Liu, Y.L., Wang, J., Li, S.Y., Li, J., and Deng, Y. (2020). Genetic analysis of oxidative and endoplasmic reticulum stress responses induced by cobalt toxicity in budding yeast. Biochim Biophys Acta Gen Subj 1864, 129516.

